# Modular evolution of secretion systems and virulence plasmids in a bacterial species complex

**DOI:** 10.1101/2021.05.20.444927

**Authors:** Lin Chou, Yu-Chen Lin, Mindia Haryono, Mary Nia M. Santos, Shu-Ting Cho, Alexandra J. Weisberg, Chih-Feng Wu, Jeff H. Chang, Erh-Min Lai, Chih-Horng Kuo

## Abstract

**Background:** Many bacterial taxa are species complexes and uncertainties regarding the organization of their genetic diversity challenge research efforts. We utilized *Agrobacterium tumefaciens*, a taxon known for its phytopathogenicity and applications in transformation, as a study system and devised strategies for investigating genome diversity and evolution of species complexes.

**Results:** We utilized 35 genome assemblies to achieve a comprehensive and balanced sampling of *A. tumefaciens*. Our confident inference of gene content and core-genome phylogeny supported a quantitative guideline for delineating 12 species and allowed for robust investigations of genes critical in fitness and ecology. For the type VI secretion system (T6SS) involved in interbacterial competition and thought to be conserved, we detected multiple losses and one horizontal gene transfer. For the tumor-inducing plasmids (pTi) and pTi-encoded type IV secretion system (T4SS) that are essential for agrobacterial phytopathogenicity, we uncovered novel diversity and hypothesized their involvement in shaping this species complex. Intriguingly, for both T6SS and T4SS, genes encoding structural components are highly conserved, whereas extensive diversity exists for genes encoding effectors and other proteins.

**Conclusions:** We demonstrated that the combination of a phylogeny-guided sampling scheme and an emphasis on high-quality assemblies provides a cost-effective approach for robust analysis in evolutionary genomics. Our strategies for multi-level investigations at scales that range from whole-genomes to intragenic domains and phylogenetic depths of between- and within-species are applicable to other bacteria. Finally, modularity observed in the molecular evolution of genes and domains is useful for inferring functional constraints and informing experimental works.

## Background

Investigation of species boundaries in bacteria is fundamentally important because species are the key identifiers for biological entities [1, 2]. However, many bacterial groups are currently unresolved and are classified as species complexes. These uncertainties regarding species boundaries hamper research, communication, and policy-making such as in healthcare guidelines, pathogen quarantine regulations, and biological resource management. Based on barriers to homologous recombination, an analysis of > 20,000 bacterial genome sequences from 91 species belonging to 13 phyla found that 21 of the previously recognized species comprise multiple biological species [3]. These 21 groups include those that are important as pathogens (e.g., *Mycobacterium tuberculosis*, *Pseudomonas aeruginosa,* and *Vibrio cholerae*) or beneficial microbes (e.g., *Lactobacillus casei* and *Sinorhizobium meliloti*). This finding highlights the ubiquity of species complexes across bacterial lineages, even for those that are extensively studied.

For such complexes, comprehensive understanding of the genetic diversity organization is required for robust species delineation, which in turn is essential for providing a reliable framework to interpretate experimental findings and to gain insights into the biology. The use of genomic information has long been suggested as a powerful approach for defining species boundaries because the comprehensive genetic information can provide definitive and potentially quantitative guidelines [4]. However, several issues regarding genomic studies of bacterial species have remained unresolved. First, while genomospecies defined by overall genome divergence were suggested to represent distinct biological entities [5–9], the exact criteria for establishing the species boundaries are disputed. Although 95% average nucleotide identity (ANI) across the conserved parts of genomes was proposed as a universal boundary for defining species in bacteria [10], this criterion was challenged [11]. Additionally, ANI values alone do not provide information such as gene content or phylogenetic relationships, which are critical in understanding biological entities [4]. Second, phylogenetic relationships among closely related bacterial strains often cannot be resolved with confidence, yet such information is fundamental for evolutionary analysis. Third, comparative genomics studies are often limited by taxon sampling and/or assembly quality of available genome sequences.

In this study, we utilized the *Agrobacterium tumefaciens* species complex, also known as *Agrobacterium* biovar 1 [12], as the study system for developing strategies that provide appropriate sampling and utilize multifaceted genomic analysis to resolve species boundaries and to investigate molecular evolution of key traits. These bacteria are known as the causative agents of crown gall disease that affects over 90 plant families [13]. More importantly, the development of *Agrobacterium*-mediated transformation has provided a critical tool for genetic manipulation in plant sciences and agricultural biotechnology [14, 15]. Due to their importance, this complex has been studied for over a century and was found to harbor extensive phenotypic and genetic diversity that continues to confound efforts to resolve their taxonomy [12–14]. Various methods, such as DNA-DNA hybridization, biochemical characteristics, and molecular markers have been used to define 10 genomospecies (i.e., G1-G9 and G13), which have continually been associated to new nomenclature, a process that causes greater confusion than resolution [5,6,16–18]. For example, the reference strain C58 used in many *A. tumefaciens* studies [15,19,20] belongs to G8, which was renamed as *Agrobacterium fabrum* in 2011 [21]. This has resulted in mixed usage of two names with different meanings (i.e., *A. tumefaciens* for the entire complex and *A. fabrum* for G8) in databases and literature. Compounding confusion, the name *Agrobacterium radiobacter* refers to *A. tumefaciens* G4 [18, 21], the entire *A. tumefaciens* complex [22], and the more divergent *Agrobacterium* biovar 2 [23]. Hereafter, we use *A. tumefaciens* in reference to the entire species complex and specific designations (i.e., G1-G9 and G13) for genomospecies.

Previous characterizations found that *A. tumefaciens* strains have multipartite genomes with one circular chromosome, one linear chromosome, and highly variable plasmids [9,23–25]. Consistent with the high levels of genetic divergence inferred from DNA-DNA hybridization [5], cross-genomospecies comparisons typically found that < 80% of the genes are conserved [7, 26]. Strikingly, > 32,000 horizontal gene transfer (HGT) events have been inferred to have shaped the evolutionary history of *A. tumefaciens* [8]. Because the HGT patterns indicated co-transfers of genes that encode coherent biochemical pathways, it was hypothesized that purifying selection on those acquired gene clusters and overall gene content drove the ecological diversification among genomospecies [8]. Moreover, the oncogenic plasmids that determine *Agrobacterium* phytopathogenicity exhibit complex modularity and transmission patterns, which further contributed to the diversification of these pathogens and their global spread [9]. However, despite the progresses, those better-characterized genomospecies (e.g., G1, G4, G7, and G8) and pathogenic strains were highly overrepresented in previous genomics studies [7–9], and such biases may affect our understanding of agrobacterial diversity and evolution.

To develop effective strategies for investigating bacterial species complexes such as *A. tumefaciens*, we started by performing targeted genome sequencing for strains in underrepresented lineages to achieve a comprehensive and balanced taxon sampling of the study system. We also limited analyses to only high-quality assemblies, which enabled detailed examinations of replicon-level synteny and confident inferences of gene presence/absence. The global view of genomic diversity and resolved phylogeny provided a robust framework for focused investigations of the genetic elements involved in key aspects of agrobacterial fitness and ecology, namely the type VI secretion system (T6SS) for interbacterial competition [7, 27] and the virulence plasmids for phytopathogenicity [13–15]. Taken together, the investigations, scaling from whole-genome, whole-replicon, gene clusters, individual genes, and intragenic protein domains, provided novel and detailed information on the evolution and genetic diversity of bacteria important in plant pathology and biotechnology. Moreover, the strategies developed in this work are applicable to the study of other bacterial species complexes.

## Results

### Genome Sampling, Molecular Phylogeny, and Divergence

Based on existing knowledge of *A. tumefaciens* diversity [5,6,16–18] and availability of genomics resources [7–9], we identified 12 strains that represent the six poorly characterized genomospecies and two strains from the sister lineage *Agrobacterium larrymoorei* to provide an ideal outgroup (Table 1). Whole-genome sequencing with substantial efforts in iterative improvements of the assemblies based on experimental and bioinformatic approaches were conducted for these 14 strains. Additionally, we conducted multiple rounds of preliminary genome-scale phylogenetic analysis to select 21 representatives from 98 *A. tumefaciens* genome assemblies available from GenBank [28] (Table 1) to yield a dataset with maximal genetic diversity without emphasis on including pathogenic strains. To ensure balanced sampling, we selected between two and five strains for each of the 10 recognized *A. tumefaciens* genomospecies. Importantly, 19 of these 35 assemblies, including nine produced in this study, are complete and most others are nearly complete (i.e., average N50 = 1.3 Mb; cf. the two chromosomes are ∼2.9 and ∼2.3 Mb, respectively).

**Table 1.** List of the genome sequences used in this study. These include 14 new genomes derived from this study and 21 additional representatives from GenBank. Two *Agrobacterium larrymoorei* strains are included as the outgroup. Species name abbreviations: *At*, *Agrobacterium tumefaciens*; *Al*, *Agrobacterium larrymoorei*.

| Species | Strain | Accession | Assembly | Coding Sequences | Pseudo-genes | Geographic Origin | Isolation Source |
| --- | --- | --- | --- | --- | --- | --- | --- |
| *At* G1 | 1D1108 | GCF_003666425 | Complete | 5,312 | 167 | MD, USA | *Euonymus* sp. |
| *At* G1 | 5A | GCF_000236125 | 50 contigs | 5,343 | 169 | MT, USA | Soil |
| *At* G1 | Ach5 | GCF_000971565 | Complete | 5,184 | 145 | CA, USA | *Achillea ptarmica* |
| *At* G1 | N2/73 | GCF_001692195 | 29 scaffolds | 5,345 | 157 | OR, USA | Cranberry |
| *At* G1 | S56 | GCF_900014385 | 6 scaffolds | 5,414 | 218 | ? | Plant |
| *At* G2 | CFBP5494 | GCF_900013495 | 5 scaffolds | 5,469 | 233 | France | Human |
| *At* G2 | CFBP5496 | GCF_005144405 | 9 contigs | 5,137 | 223 | France | Human |
| *At* G2 | CFBP5875 | GCF_005221365 | Complete | 4,570 | 146 | Belgium | Ditch water |
| *At* G3 | CFBP6623 | GCF_005221385 | Complete | 5,081 | 180 | France | Antiseptic flask |
| *At* G3 | CFBP6624 | GCF_005221425 | Complete | 5,157 | 148 | France | Human |
| *At* G4 | 183 | GCF_004023565 | Complete | 5,051 | 199 | Tunisia | *Prunus dulcis* |
| *At* G4 | 186 | GCF_002591665 | Complete | 5,298 | 207 | CA, USA | *Juglans regia* |
| *At* G4 | 12D1 | GCF_003667905 | Complete | 5,005 | 167 | ? | ? |
| *At* G4 | 1D1460 | GCF_003666445 | Complete | 5,290 | 269 | CA, USA | *Rubus* sp. |
| *At* G5 | CFBP6625 | GCF_005221465 | Complete | 5,446 | 198 | France | Food |
| *At* G5 | CFBP6626 | GCF_005221445 | Complete | 5,080 | 199 | France | Human |
| *At* G5 | F2 | GCF_000219665 | 8 contigs | 5,085 | 135 | Harbin, China | Soil |
| *At* G6 | CFBP5499 | GCF_005221325 | Complete | 5,654 | 240 | South Africa | *Dahlia* sp. |
| *At* G6 | CFBP5877 | GCF_005221345 | Complete | 5,463 | 236 | Israel | *Dahlia* sp. |
| At G7 | 1D1609 | GCF_002943835 | Complete | 5,539 | 254 | CA, USA | *Medicago sativa* |
| At G7 | CFBP4996 | GCF_005144435 | 10 contigs | 5,741 | 213 | UK | *Flacourtia ramontchi* |
| At G7 | CFBP7129 | GCF_005221405 | Complete | 5,927 | 285 | Tunisia | *Pyrus communis* |
| At G8 | 12D13 | GCF_003667945 | Complete | 5,083 | 178 | ? | ? |
| At G8 | 1D132 | GCF_003667725 | Complete | 5,176 | 169 | CA, USA | *Cerasus pseudocerasus* |
| At G8 | ATCC31749 | GCF_002916755 | 4 contigs | 4,560 | 811 | China | Plant |
| At G8 | C58 | GCF_000092025 | Complete | 5,355 | 29 | NY, USA | *Cerasus pseudocerasus* |
| At G9 | CFBP5506 | GCF_005144495 | 15 contigs | 4,236 | 157 | Australia | Soil |
| At G9 | CFBP5507 | GCF_005144505 | 16 contigs | 5,299 | 253 | Australia | Soil |
| At G9 | GBBC3283 | GCF_007002765 | 48 contigs | 4,946 | 203 | Belgium | *Solanum lycopersicum* |
| At G13 | CFBP6927 | GCF_900012615 | 9 scaffolds | 4,722 | 94 | France | Rhizospheric soil from *Prunus persicae* |
| At G13 | S2 | GCF_000723345 | 60 contigs | 5,574 | 167 | ? | ? |
| At G14 | KCJK1736 | GCF_001641425 | 41 contigs | 4,933 | 162 | FL, USA | *Bos taurus* feces |
| At G15 | MAFF210266 | GCF_007002865 | 26 contigs | 5,199 | 162 | Japan | *Cucumis melo* |
| AI | CFBP5473 | GCF_005145045 | Complete | 4,777 | 142 | FL, USA | *Ficus benjamina* |
| AI | CFBP5477 | GCF_005144425 | 24 contigs | 4,657 | 95 | Italy | ? |

Based on the comprehensive sampling of *A. tumefaciens* diversity, we identified a core genome of 2,093 single-copy genes, which correspond to ∼40% of the genes annotated in each individual genome sequence. Compared to previous studies that conducted genome-based phylogenetic analysis for *Agrobacterium* or higher taxonomic ranks [8,9,29], the more focused sampling in this study yielded a higher core gene count by one-to-two orders of magnitude. This increase in core gene count and the improvement in taxon sampling allowed for the inference of a well-resolved maximum likelihood phylogeny of the *A. tumefaciens* species complex (Figure 1A). Each of the 10 currently recognized genomospecies forms a distinct monophyletic clade with > 80% bootstrap support. Additionally, we identified two novel genomospecies, G14 and G15, each represented by a single strain. The pattern of overall genome similarities exhibits a discrete multimodal distribution that supports use of a 95% ANI cutoff for delineating bacterial species [10] and quantifies the divergence of the *A. tumefaciens* complex from its most closely related sister lineage (Figure 1B).

**Figure 1.**
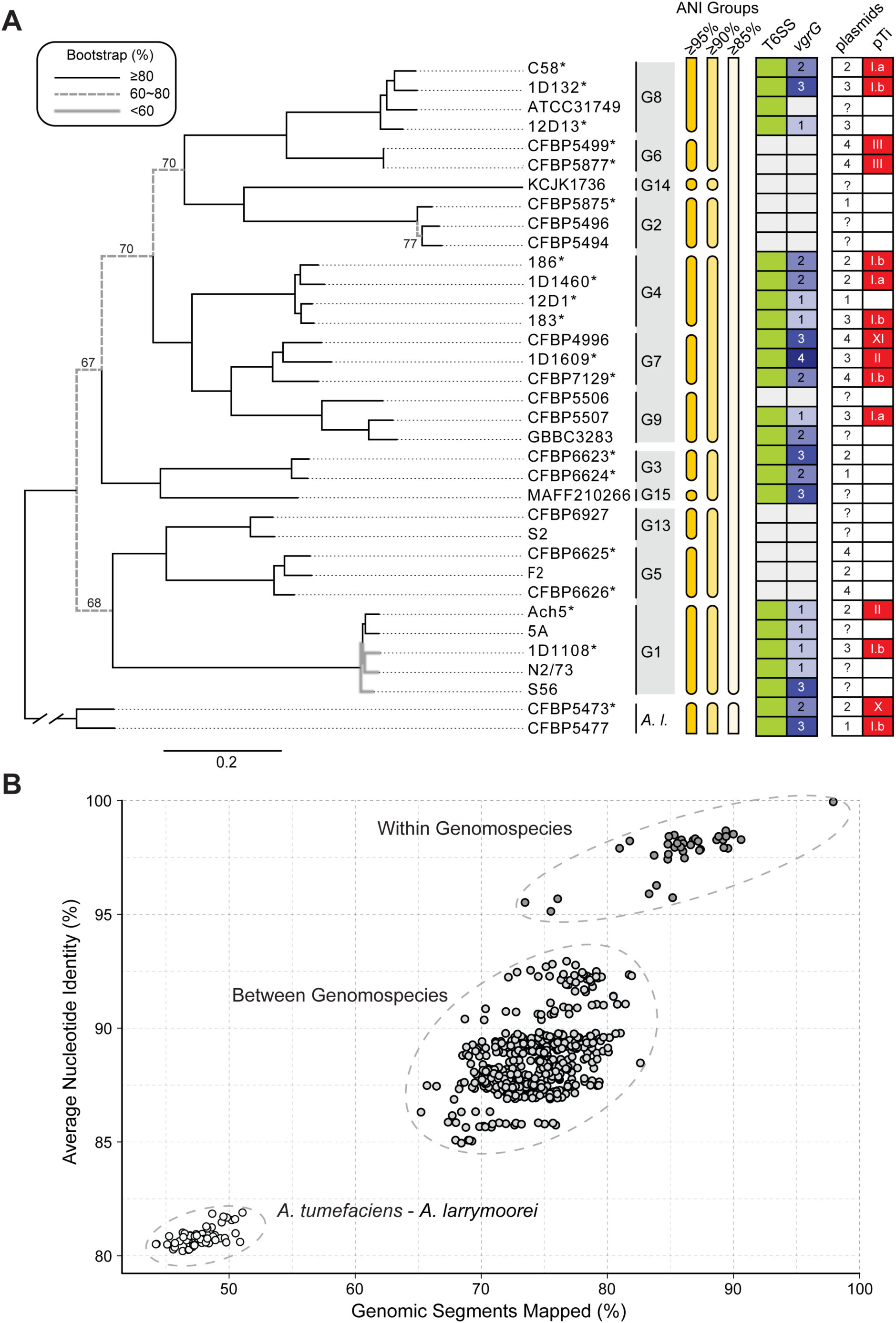
Relationships among representatives of the *Agrobacterium tumefaciens* species complex. The sister species *Agrobacterium larrymoorei* (*A. l.*) is included as the outgroup. (A) Maximum likelihood phylogeny based on a concatenated alignment of 2,093 single-copy genes shared by all 35 genomes (635,594 aligned amino acid sites). Bootstrap support values in the range of 60-80% are labeled. Strains with complete genome assemblies are highlighted with an asterisk (‘*’). The genomospecies assignments (i.e., G1-G9 and G13-G15) are labeled to the right of strain names. The three *A. tumefaciens* supergroups are indicated by the colored background of the genomospecies assignments. Information to the right of the genomospecies assignments shows the grouping of genomes according to different cutoff values of genome-wide average nucleotide identity (ANI), the presence/absence of type VI secretion system (T6SS)-encoding gene cluster (green: present; white: absent), copy number of *vgrG* (white background: absent), number of plasmids, and the tumor-inducing plasmid (pTi) type based on k-mer profile (white background: absent). (B) Pairwise genome similarities based on the percentages of genomic segments mapped and the ANI values.

The 12 *A. tumefaciens* genomospecies were classified into seven groups based on 90% ANI, and further assigned to three supergroups according to the phylogeny (Figure 1A). Based on a time-calibrated phylogeny [9], the most recent common ancestor (MRCA) of *A. tumefaciens* emerged ∼48 million years ago (Mya), the three supergroups emerged ∼40 Mya, the seven groups emerged ∼30 Mya, and most of the recognized genomospecies emerged ∼2-10 Mya. The inferred rapid radiation in the early history of *A. tumefaciens* likely prevented confident resolution of those deeper relationships in previous studies [8,9,29]. Nevertheless, with improvements in the taxon sampling of this work, we observed ∼70% bootstrap support for those early nodes (Figure 1A). This fully-resolved organismal tree provides a strong framework for our downstream examination of gene and domain phylogenies.

The high levels of assembly completeness provided confident inference of gene content comparisons. The principal coordinate analysis and hierarchical clustering results indicated that all 12 *A. tumefaciens* genomospecies are similar to one another while distinct from *A. larrymoorei* (Supplementary Figure S1A and S1C). Nonetheless, with the exception of G4 and G7, these genomospecies are distinguishable based on gene content (Supplementary Figure S1B and S1D). This finding suggests that despite the extensive HGT inferred within this complex [8], the genomospecies defined by 95% ANI likely represent distinct biological entities.

### Diversity and Evolution of the T6SS Genes

The fully-resolved organismal phylogeny and confidence in gene content inference, particularly for the cases of gene absence, provide a robust framework for evolutionary analysis of key traits. We first focus on genes encoding the T6SS, a phage tail-like contractile nanomachine commonly found in Proteobacteria and used to inject effectors into eukaryotic or bacterial cells. The T6SS has major roles in pathogenesis, symbiosis, and interbacterial competition [30–33]. For *A. tumefaciens*, the T6SS is a key weapon for *in planta* competition between different genomospecies [7] and against other bacteria [27]. Thus, investigating the diversity and evolution of T6SS genes may shed light on a trait that influences the ecology and evolution of *A. tumefaciens*.

In a previous study that examined four *A. tumefaciens* genomospecies, T6SS-mediated anti-bacterial activity was observed for all 11 strains sampled and thought to be a conserved trait of this species complex [7]. To our surprise, among the 33 *A. tumefaciens* strains examined in this work, a patchy distribution of the T6SS genes was observed (Figure 1). Gene absences are in strains corresponding to previously under-characterized genomospecies and were confirmed by examining syntenic regions and using TBLASTN [34] to search entire genome sequences. For strains encoding a T6SS, corresponding genes are consistently located on the linear chromosome and mostly form a cluster of ∼20 genes organized as two adjacent and oppositely oriented *imp* and *hcp* operons [7, 35] (Figure 2). Some strains harbor accessary loci containing *vgrG* (involved in effector delivery) and other T6SS genes located elsewhere on the linear chromosome [7,33,35] (Supplementary Figure S2). The T6SS gene phylogeny is largely congruent with the species tree (Figure 2). One notable exception is that the MRCA of G1 appears to have acquired the T6SS genes from a G8-related donor. Consistent with this inference, the T6SS genes in G1 strains are located in a different chromosomal location compared to strains of other genomospecies (Supplementary Figure S2). Based on these observations, it is likely that a T6SS gene cluster was present in the MRCA of G8-G6-G14-G2-G4-G7-G9-G3-G15 and at least two independent losses have occurred in G6 and G14-G2. Regarding the ancestral state in the MRCA of the *A. tumefaciens* complex, presence of the T6SS genes appears to be a more parsimonious hypothesis based on the presence of these genes in the outgroup (Figure 2). However, the lack of synteny conservation between the linear chromosomes of *A. tumefaciens* and *A. larrymoorei* and the variable locations of *vgrG* homologs (Supplementary Figure S2) suggest that multiple independent origins are also possible. For broader scales, the T6SS genes have a patchy distribution among Rhizobiaceae [33, 36], indicating that these genes are not essential for these bacteria and have high rates of gains and losses. Consistent with this, multiple pseudogenes confirmed by manual curation of annotation were found (e.g., *tssH* in S56 and *tssA*/*vgrG* in ATCC31749) (Figure 2), suggesting that for some strains these gene clusters are in the process of degradation and will be eventually lost.

**Figure 2.**
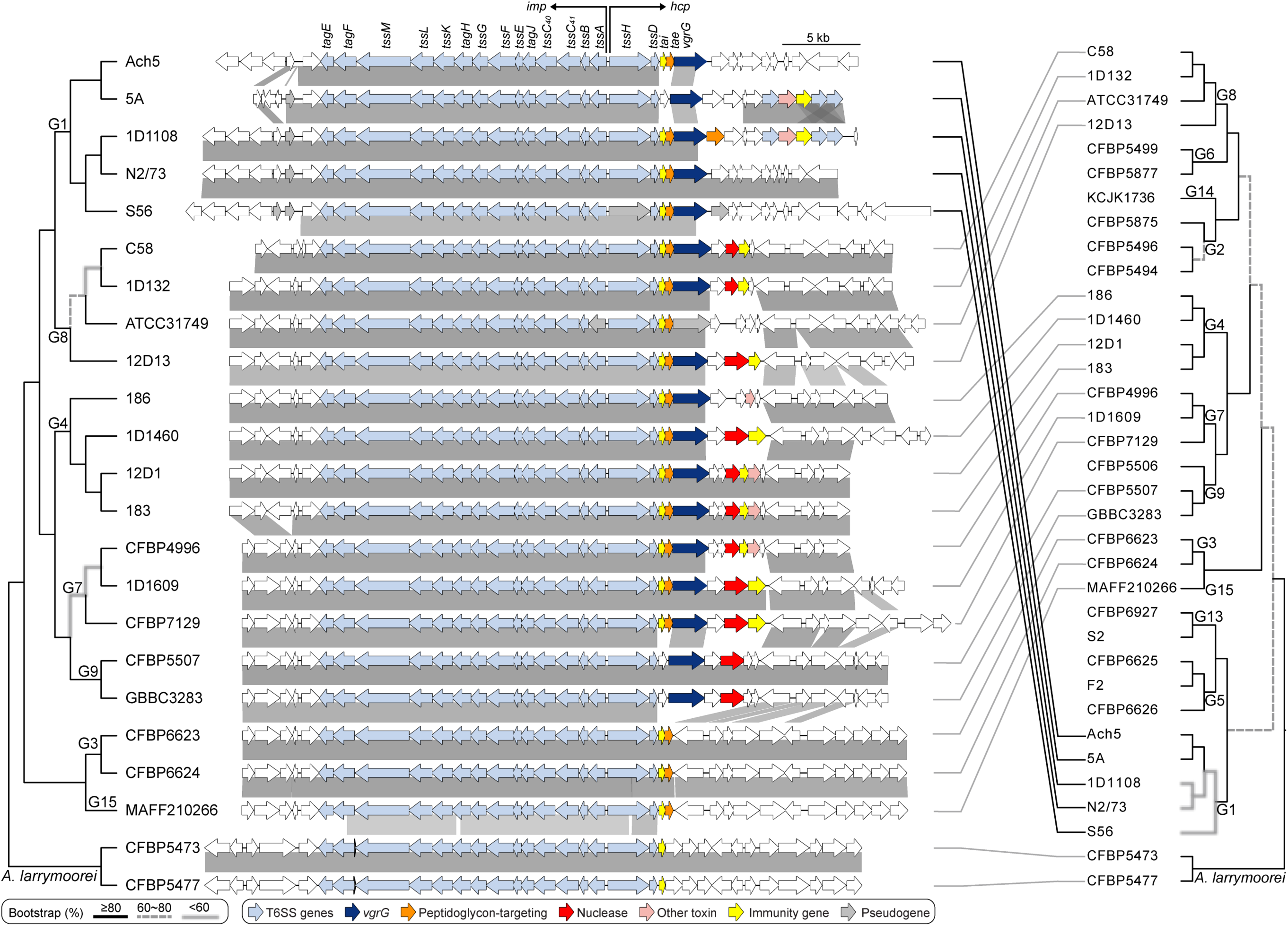
Phylogeny and organization of the T6SS gene cluster. The maximum likelihood phylogeny was inferred based on a concatenated alignment (5,960 aligned amino acid sites) of 14 core T6SS genes, including *tagE*, *tagF*, *tssM*, *tssL*, *tssK*, *tagH*, *tssG*, *tssF*, *tssE*, *tagJ*, *tssC40*, *tssC41*, *tssB*, and *tssD*. Two other core genes, *tssA* and *tssH*, are excluded because some homologs are pseudogenized. Genes downstream of *tssD* (e.g., *tai*, *tae*, and *vgrG*) are excluded due to variable presence. The species phylogeny on the right is based on Figure 1. Genes are color-coded according to annotation, syntenic regions are indicated by gray blocks.

Examining synteny revealed that the *imp* operon, which encodes the majority of T6SS structural components [37], is more conserved in gene composition and order than the *hcp* operon, which often has different genes downstream of *vgrG* (Figure 2). This genetic diversity may play a key role in interbacterial competition because genes downstream of *vgrG* include those that encode effector and immunity (EI) protein pairs [7]. The agrobacterial T6SS effectors often correspond to different toxins and the cognate immunity proteins provide protection against self-intoxication [7,27,33,36]. The rapid evolution of EI gene pairs is illustrated by three examples. First, despite the low levels of sequence divergence among G1 strains (i.e., > 98% ANI in all pairwise comparisons), different genes are found downstream of their *vgrG* homologs and this variation is not consistent with either the species phylogeny or the T6SS core gene phylogeny. Second, strain CFBP4996 of G7 has homologs of the same EI gene pair as strains 12D1 and 183 of G4, rather than with other members of G7. Third, in both G3 strains, *vgrG* and the associated EI genes are located elsewhere on the linear chromosome, rather than being a part of the *hcp* operon (Figure 2 and Supplementary Figure S2). These results suggest that recombination involving gene modules has contributed to the genetic diversity of these *A. tumefaciens* T6SS EI gene pairs.

### Modularity of VgrG and Its Associated EI Pairs

The knowledge that *vgrG* homologs encode proteins with distinct C-terminal domains responsible for binding specificities of different T6SS effectors for delivery suggested that each *vgrG* homolog and its downstream EI gene pair may evolve as a functional module [33, 36]. Here, we sought to investigate the patterns of gene co-occurrence and intra-module recombination to better understand the diversity and evolution of these genes. For in-depth investigation of *vgrG* evolution, we began by examining domain architecture of VgrG proteins and uncovered eight distinct domains (Supplementary Figure S3). Based on differences in domain composition, the 44 *vgrG* homologs, including 17 associated with the main T6SS gene cluster and 27 associated with accessory loci, were classified into six major types and nine subtypes (Figure 3). Only three of the domains are present in all VgrG variants. The N-terminal domain 1 is the most conserved (Supplementary Figure S3) and the only one found in databases. This domain corresponds to TIGR03361, which accounts for ∼66-77% of the protein length and the bulk of the structures that forms a trimeric complex analogous to a phage tail spike [38, 39] (Supplementary Figure S4). For domain 5 that was found in all *vgrG* homologs belonging to subtypes A1-A3 and E1 (Figure 3), the presence of this domain is perfectly correlated with the presence of a downstream DUF4123-domain-containing gene (Figure 4). Because this DUF4123 domain acts as an adaptor/chaperone for effector loading onto VgrG in *A. tumefaciens* [36] and *Vibrio cholerae* [40, 41], this strong co-occurrence suggests specific interactions between VgrG domain 5 and DUF4123. Thus, combining domain analysis with gene co-occurrence provides a new strategy for predicting the interacting domains of VgrG with other T6SS components.

**Figure 3.**
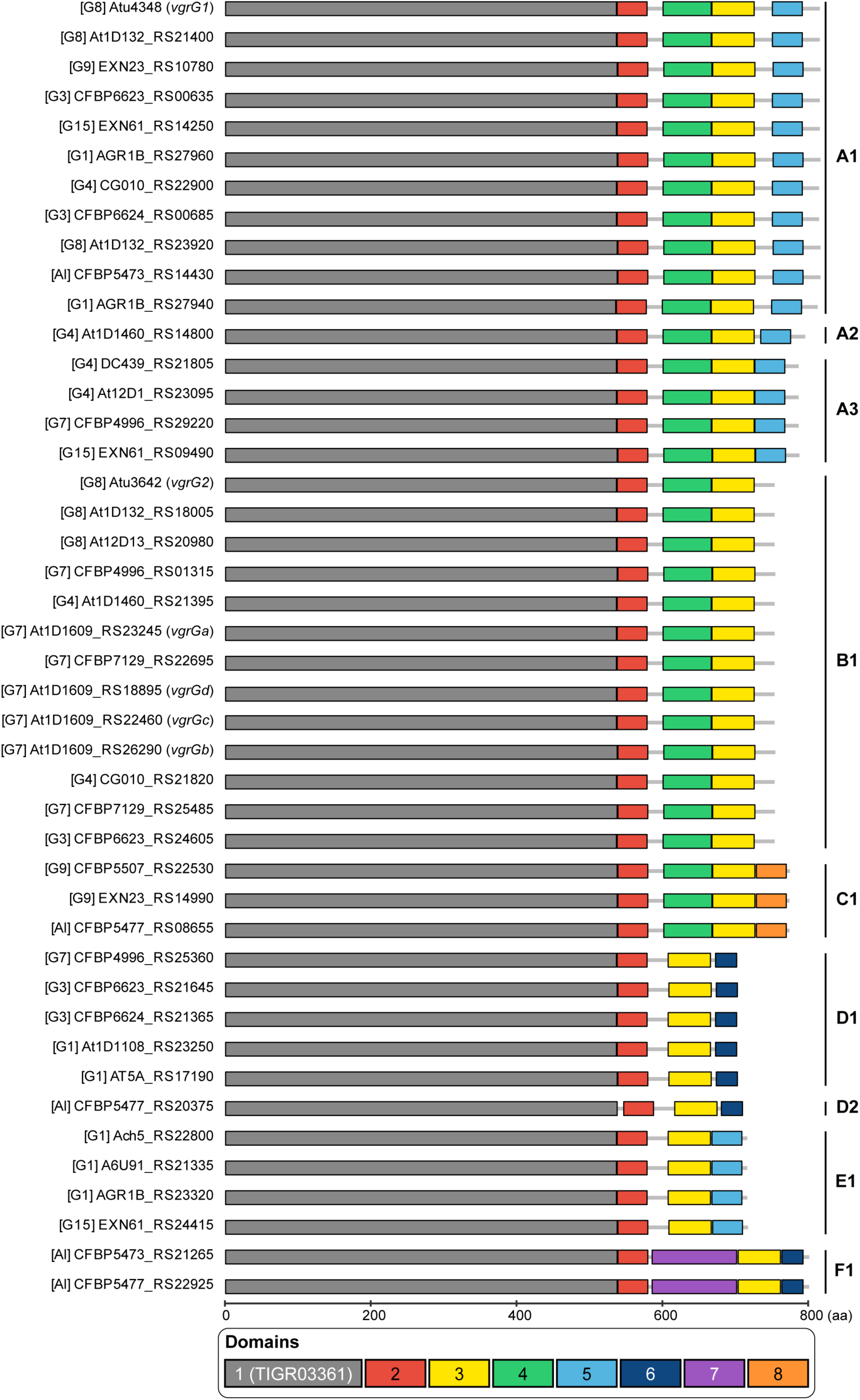
Domain organization and classification of *vgrG* homologs. Six major types (i.e., A-F) and nine subtypes were identified and labeled on the right. For each homolog, the genomospecies assignment is provided in square brackets, followed by the locus tag. The gene names are provided in parenthesis for those functionally characterized homologs (i.e., *vgrG1-2* for C58 homologs and *vgrGa-d* for 1D1609 homologs).

**Figure 4.**
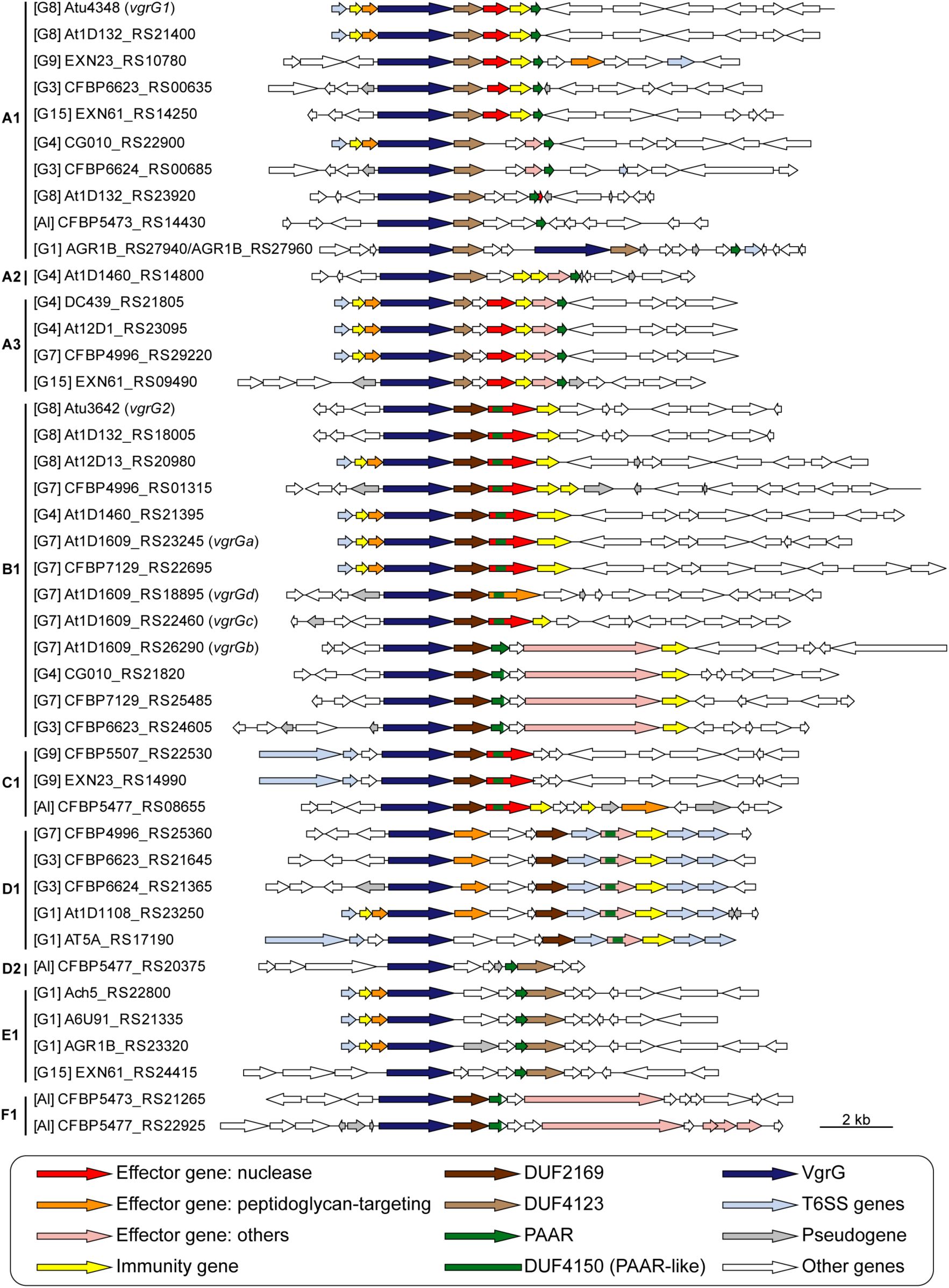
Gene neighborhoods of *vgrG* homologs. The grouping and labeling of *vgrG* homologs are based on the convention used in Figure 3. For each *vgrG* homolog, three upstream genes are plotted to illustrate if it is associated with the main T6SS gene cluster or not, and 10 downstream genes are plotted to illustrate putative effector/immunity genes. Two A1-type homologs from the strain S56 (i.e., AGR1B_RS27940 and AGR1B_RS27960) are in close association with each other and plotted together.

Intriguingly, despite conservation of domain architecture within each subtype (Figure 3), the phylogenies inferred from the three domains encoded by all *vgrG* homologs do not have the same topology (Supplementary Figure S5). For domain 1, sequences from the same subtype do not always form monophyletic groups. For domains 2 and 3, the short sequence lengths limited the phylogenetic resolution; nonetheless, low divergence within the same subtype and high divergence between subtypes were observed. These patterns suggest that each domain-encoding region evolved independently and can recombine between subtypes.

At the level of gene cluster organization, *vgrG* homologs within a subtype can have distinct downstream genes (e.g., A1 and B1), regardless of whether they are associated with the main T6SS gene cluster (Figure 4 and Supplementary Dataset S1A). These findings suggest that in addition to domain shuffling among *vgrG* homologs, recombination also facilitated novel *vgrG*-effector pairings in the evolution of these T6SS genes.

When T6SS diversity was examined in a phylogenetic context, numbers and types of *vgrG* homologs, as well as their linked EI genes, lack strong correlations with species phylogeny (Figure 5). Based on our manual curation of *vgrG*-associated genes, a total of 63 putative effector genes were identified (Supplementary Dataset S1A). Among these, peptidoglycan-targeting toxins and nucleases are the two most commonly found categories with 21 each. This finding is consistent with an investigation of > 1,000 T6SS effectors sampled from 466 species in the phylum Proteobacteria [32], which also found these two as the dominant categories and suggested that they play complementary roles in T6SS-mediated competition.

**Figure 5.**
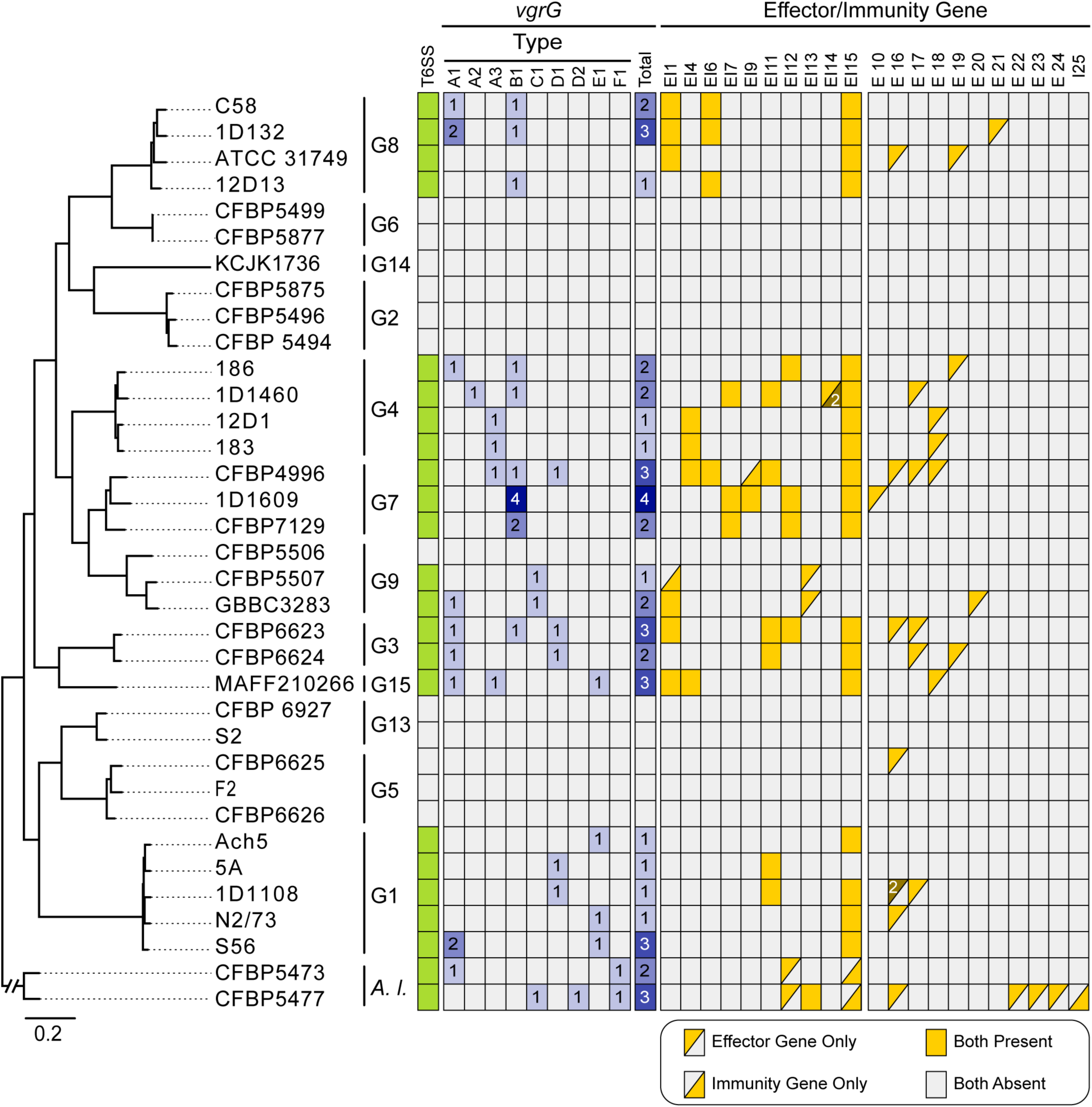
Phylogenetic distribution of *vgrG* homologs and T6SS effector/immunity genes. The species tree is based on Figure 1. Gene presence is illustrated with colored cells in the heatmap, gene copy numbers are labeled when applicable.

### The Virulence Plasmids and Associated Genes

The tumor-inducing plasmids (pTi) are an important component of *A. tumefaciens* genomes. These large accessory replicons harbor the virulence (*vir*) regulon genes that encode the Vir proteins and type IV secretion system (T4SS) for processing and delivering a transfer DNA (T-DNA) into plant cells, and are essential for agrobacterial phytopathogenicity [13–15]. Among the 35 strains examined, we identified 15 pTi sequences (Table 2). Two novel putative pTi (i.e., pTiCFBP4996 and pTiCFBP5473) were found in the 14 newly sequenced strains. This efficiency of discovering novel pTi types is surprising, given our previous study that defined pTi types I-VI was based on comprehensive sampling of diverse historical collections containing 162 *Agrobacterium* strains [9]. This finding demonstrates the importance and usefulness of a phylogeny-guided approach for investigating genetic diversity. Our recent examination of > 4,000 Rhizobiaceae plasmids assigned these two novel pTi to types X and XI, and found that both types are species-specific [42], These two novel pTi are distinctive in their large sizes (Table 2), gene organization (Figure 6), gene content (Supplementary Figure S6), and sequence divergence of core genes (Supplementary Figure S7). Moreover, their T-DNA regions also differ from the typical sizes of ∼18-26 kb observed in types I-III pTi (Figure 6). For the tumorigenic strain CFBP5473 (Supplementary Figure S8), the predicted T-DNA border sequences flank an extraordinarily large (∼93 kb) region that contains all of the *vir* regulon genes in addition to the typical T-DNA-associated genes (e.g., synthesis of opine and plant hormone) (Figure 6). This reflects either a translocation of a T-DNA border sequence or reliance on a non-canonical T-DNA border sequence that we could not identify. For pTiCFBP4996, its 7-kb T-DNA is predicted to contain only four genes (i.e., two correspond to opine synthesis and two encode hypothetical proteins). Plant hormone synthesis genes, which are necessary to cause visible disease symptoms, were not identified in this predicted T-DNA region or elsewhere on this plasmid. Consistent with predictions, strain CFBP4996 did not induce tumor formation when inoculated onto stems of tomato plants (Supplementary Figure S8). Considering that pTiCFBP4996 encodes all essential *vir* genes required for T-DNA processing and transfer, CFBP4996 may serve as a naturally disarmed strain capable of T-DNA transfer without causing diseases.

**Figure 6.**
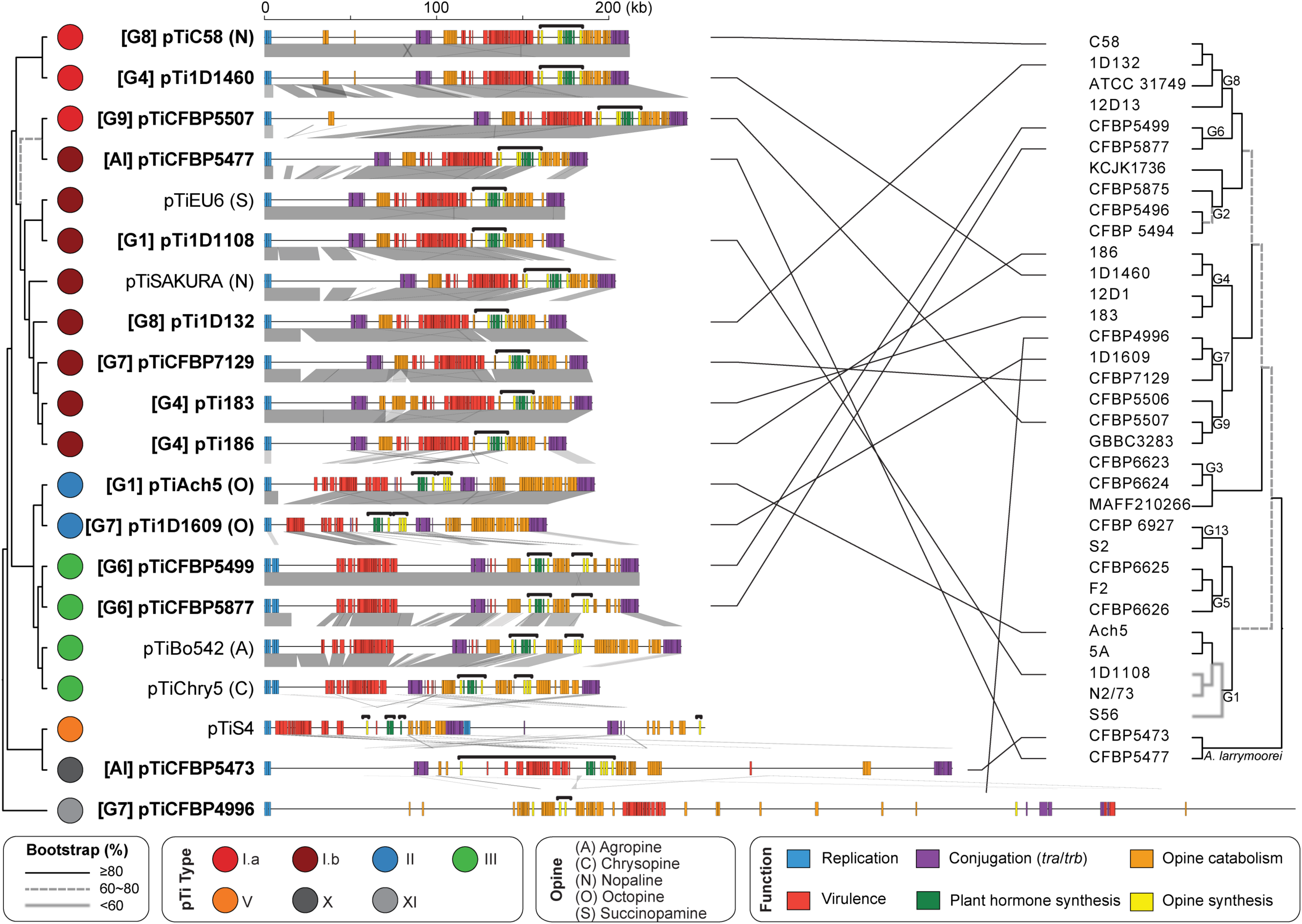
Molecular phylogeny and global alignment of pTi. The maximum likelihood phylogeny was inferred based on the concatenated alignment of 21 core genes and 8,534 aligned amino acid sites (**Supplementary Figure S6A**). The species phylogeny on the right is based on Figure 1. The pTi sequences derived from this study are highlighted in bold. For those with relevant information available, the genomospecies assignments are indicated in square brackets and the opine types are indicated in parentheses. For the alignment, all plasmids are visualized in linear form starting from the replication genes. Other gene clusters are color-coded according to functions. Syntenic regions are indicated by gray blocks.

**Table 2.** List of the pTi sequences analyzed. These include 15 from the genome data set listed in Table 1 and five additional representatives downloaded from GenBank. The pTi type assignments are based on k-mer profiles. A type V pTi from *Agrobacterium vitis* is included as the outgroup. Species name abbreviations: *At*, *Agrobacterium tumefaciens*; *Al*, *Agrobacterium larrymoorei*; *Av*, *Agrobacterium vitis*.

| Species | Strain | pTi | pTi Type | Opine Type | Accession | Size (bp) | Coding Sequences | Geographic Origin | Isolation Source |
| --- | --- | --- | --- | --- | --- | --- | --- | --- | --- |
| *At* G1 | 1D1108 | pTi1D1108 | I.b | ? | NZ_CP032925 | 176,213 | 159 | MD, USA | *Euonymus* sp. |
| *At* G1 | Ach5 | pTiAch5 | II | Octopine | NZ_CP011249 | 194,264 | 153 | CA, USA | *Achillea ptarmica* |
| *At* G1 | 183 | pTi183 | I.b | ? | NZ_CP029048 | 192,674 | 173 | Tunisia | *Prunus dulcis* |
| *At* G4 | 186 | pTi186 | I.b | ? | NZ_CP042277 | 177,577 | 159 | CA, USA | *Juglans regia* |
| *At* G4 | 1D1460 | pTi1D1460 | I.a | ? | NZ_CP032929 | 214,233 | 198 | CA, USA | *Rubus* sp. |
| *At* G4 | MAFF301001 | pTiSAKURA | I.b | Nopaline | NC_002147 | 206,479 | 195 | Japan | *Cerasus pseudocerasus* |
| *At* G6 | CFBP5499 | pTiCFBP5499 | III | ? | NZ_CP039893 | 220,025 | 178 | South Africa | *Dahlia* sp. |
| *At* G6 | CFBP5877 | pTiCFBP5877 | III | ? | NZ_CP039902 | 220,025 | 181 | Israel | *Dahlia* sp. |
| *At* G7 | 1D1609 | pTi1D1609 | II | Octopine | NZ_CP026926 | 166,117 | 138 | CA, USA | *Medicago sativa* |
| *At* G7 | CFBP4996 | pTiCFBP4996 | XI | ? | NZ_CM016551 | 605,495 | 521 | UK | *Flacourtia ramontchi* |
| *At* G7 | CFBP7129 | pTiCFBP7129 | I.b | ? | NZ_CP039927 | 189,955 | 176 | Tunisia | *Pyrus communis* |
| *At* G8 | 1D132 | pTi1D132 | I.b | ? | NZ_CP033026 | 177,577 | 160 | CA, USA | *Cerasus pseudocerasus* |
| *At* G8 | C58 | pTiC58 | I.a | Nopaline | NC_003065 | 214,233 | 197 | NY, USA | *Cerasus pseudocerasus* |
| *At* G9 | CFBP5507 | pTiCFBP5507 | I.a | ? | NZ_CM016546 | 248,634 | 225 | Australia | Soil |
| ? | Bo542 | pTiBo542 | III | Agropine | NC_010929 | 244,978 | 223 | Germany | *Dahlia* sp. |
| ? | Chry5 | pTiChry5 | III | Chrysopine | KX388536 | 197,268 | 210 | FL, USA | *Chrysanthemum*<br>sp. |
| ? | EU6 | pTiEU6 | I.b | Succinamopine | KX388535 | 176,375 | 194 | CT, USA | *Euonymus* sp. |
| Al | CFBP5473 | pTiCFBP5473 | X | ? | NZ_CP039694 | 404,101 | 360 | FL, USA | *Ficus benjamina* |
| Al | CFBP5477 | pTiCFBP5477 | I.b | ? | NZ_CM016547 | 190,036 | 169 | Italy | ? |
| Av | S4 | pTiS4 | V | Vitopine | NC_011982 | 258,824 | 193 | Hungary | *Vitis* sp. |

For replicon-level comparisons, types II and III pTi are more similar to each other than to type I pTi based on gene content (Supplementary Figure S6) and core gene phylogeny (Supplementary Figure S7). Within type I, the two subtypes (i.e., I.a and I.b) are distinguishable by gene content (Supplementary Figure S6) but do not form mutually exclusive clades in the core-gene phylogeny (Supplementary Figure S7). All putative pTi, including the two novel types and pTiS4 (i.e., type V from distantly-related *Agrobacterium vitis*), contain genes for the T4SS that mediates T-DNA transfer into plant cells (i.e., *virB1*-*B11* and *virD4*) and the corresponding two-component regulatory system (i.e., *virA* and *virG*) (Figure 7). Strain 1D1609 is a notable case because its *virA* and *virJ* are located on another plasmid, rather than the pTi [26]. For the other *vir* regulon genes, several differences among pTi types were observed (Figure 7). For example, while all pTi harbor a conserved *virE3a* that facilitates T-DNA protection and entry into host [43], type I pTi harbor one or two additional copies of *virE3* that belong to different sequence types (i.e., sharing the same annotation but classified as distinct homologs due to sequence divergence). Similarly, *virF*, which encodes an F-box protein that is a putative host-range determinant [44], can be classified into three sequence types with distinct distributions. Those less well-characterized *vir* genes, such as *virD3* [45, 46], *virJ* [47], and *virP*, are also distributed differently among pTi types. Finally, in addition to the presence/absence of individual genes, the overall organization of *vir* regulons also differ among these pTi (Supplementary Figure S9). All type I pTi are conserved in sharing a ∼40 kb region that contains all *vir* genes. In comparison, locations of *vir* genes are more variable among type II/III pTi; *virF/P* (and *virQ/H* if present) are located ∼5-50 kb away from the main *vir* gene cluster. Other than *vir* genes, the gene content and organization of T-DNA are also different (Figure 8). All type I pTi have one single T-DNA region, while types II and III have two and type V have four, respectively [9] (Figure 6). Within the T-DNA regions, the plant hormone synthesis genes (i.e., *tms1*/*iaaM*, *tms2/iaaH*, and *ipt*) are the most conserved ones, while others are more variable (Figure 8). Taken together, this genetic variation may contribute to the host range differences observed among strains harboring different types of pTi [48].

**Figure 7.**
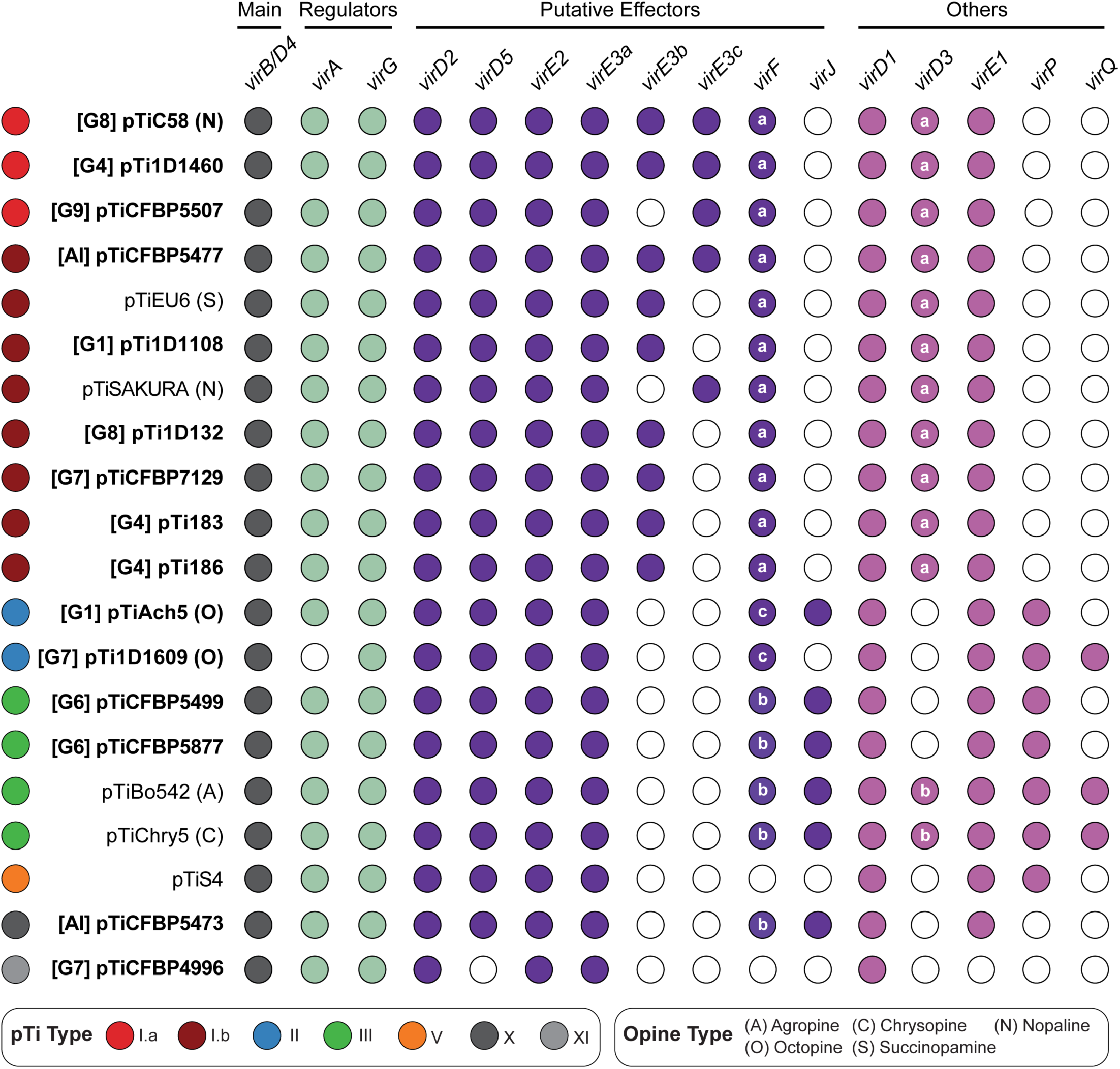
Distribution of key *vir* genes among the putative pTi. The pTi sequences derived from this study are highlighted in bold. For those with relevant information available, the genomospecies assignments are indicated in square brackets and the opine types are indicated in parentheses. Gene presence and absence are indicated by filled and empty circles, respectively. The main *vir* genes include the components of the type IV secretion system (T4SS; *virB1*-*B11* and *virD4*). For *virE3*, the three sequence types are listed separately. For *virF* and *virD3*, the sequence types are labeled inside the circles. For 1D1609, the genes *virA* and *virJ* are located on another plasmid and plotted as absent in this figure. The locus tags are provided in **Supplementary Dataset S1B**.

**Figure 8.**
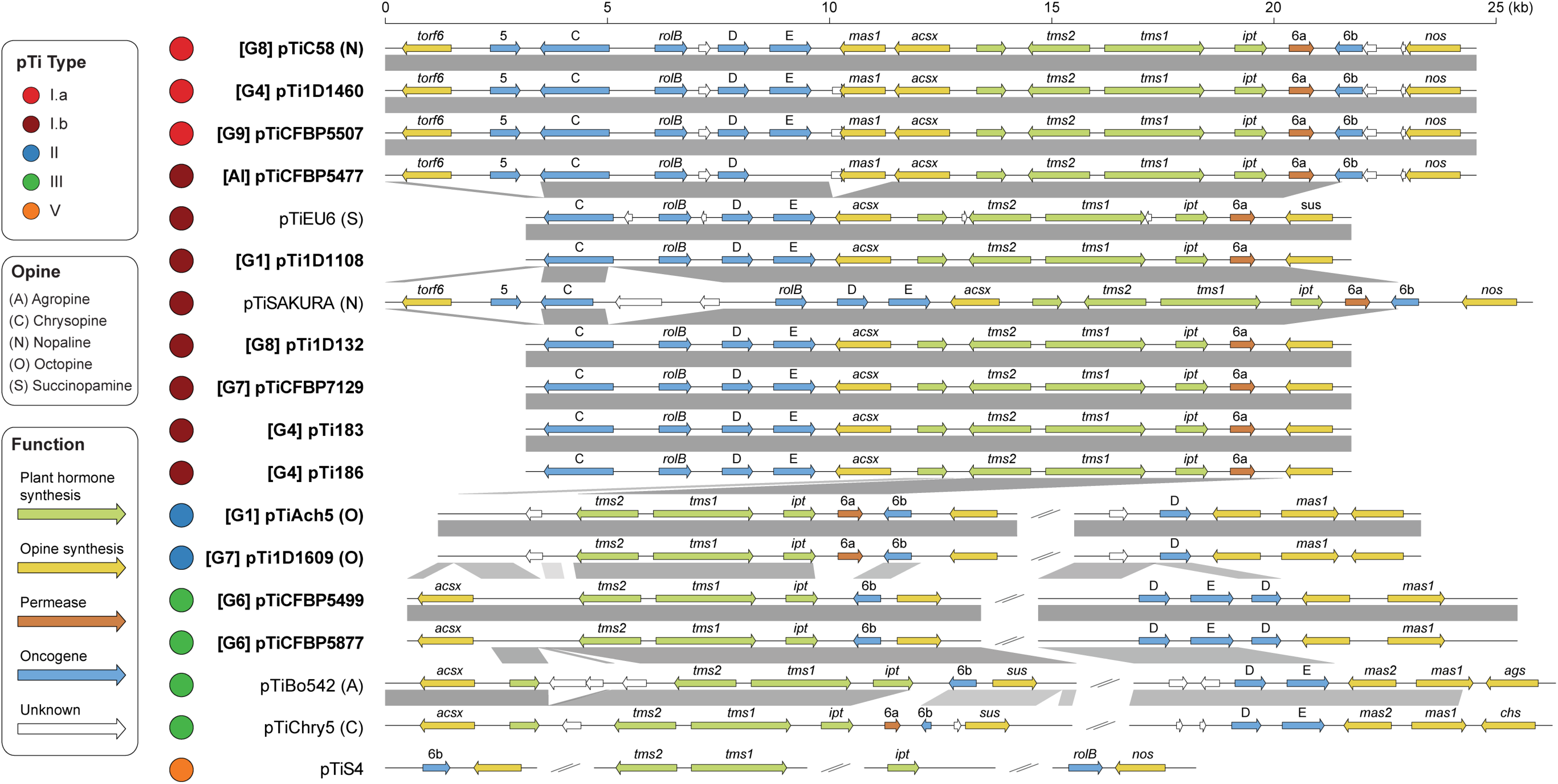
Organization of the transfer DNA (T-DNA) on tumor-inducing plasmids (pTi). Two unusual putative pTi sequences (i.e., pTiCFBP4996 and pTiCFBP4573) are excluded, other pTi sequences derived from this study are highlighted in bold. For those with relevant information available, the genomospecies assignments are indicated in square brackets and the opine types are indicated in parentheses. Genes are color-coded according to annotation, syntenic regions are indicated by gray blocks.

## Discussion

### Biological Entities at the Species Level and Above

Based on the divergence of core gene sequences (Figure 1A) and gene content (Supplementary Figure S1), 95% ANI is a reliable approach for defining species within the *A. tumefaciens* complex, as is the case for most other bacteria [10]. The discrete multimodal distribution of genome similarities (Figure 1B) suggested that there are genetic barriers between different genomospecies, which may be explained by neutral processes and/or selection [8, 10]. Regardless of the exact mechanisms, these patterns supported application of the biological species concept, which is based on genetic barriers, to these *A. tumefaciens* genomospecies and other bacteria [3]. With the continuing drop in sequencing cost, it is foreseeable that ANI analysis can serve as a practical or even a standard approach for accurate classification of additional strains, which in turn could facilitate research and communication, and ideally leads to improvements in bacterial taxonomy for basic works and applications.

At the above-species level, *A. tumefaciens* genomospecies exhibited some intriguing patterns of genome divergence. In a previous study that compared ∼90,000 prokaryotic genomes, it was extremely rare to find ANI values in the range of 82-96% [10]. In other words, strains either belong to the same natural biological entity at the species level and have > 95% ANI, or belong to different species and have < 82% ANI. This observation could be due to biases in the sampling of available genomes, or the all-against-all pairwise comparisons included mostly distantly-related species [11]. In our study that provided a detailed examination of closely-related species, the ∼85-93% ANI among *A. tumefaciens* genomospecies (Figure 1B) indicated that the *A. tumefaciens* complex is indeed a coherent entity with high divergence from its closest sister lineage within the same genus (i.e., *A. larrymoorei*). The driving forces for maintaining species complexes and the prevalence of such above-species level entities are interesting questions that require further investigations. For the *A. tumefaciens* complex, although the nomenclature originated from its phytopathogenicity, it is well-established that this group contains both pathogenic and non-pathogenic strains that differ in the possession of an oncogenic plasmid (i.e., pTi) or not [12]. The promiscuous nature of their pTi [9,49–51] (Figure 6) suggested that lineages within this complex may experience frequent transitions between pathogenic and non-pathogenic lifestyles, and such shared ecological niches may be the force that maintains the coherence of this species complex. Compared to sister lineages (e.g., *A. larrymoorei* and *A. rubi*), the more diverse host range of *A. tumefaciens* [12, 13] may have facilitated the divergence of this complex into multiple genomospecies. To test this hypothesis, better sampling of these sister lineages is required.

At the genus level and above, genome-based classification is more challenging. The ANI approach is expected to have limited resolution when nucleotide sequence identities are below ∼80% [10]. Moreover, the fractions of genome sequences alignable for ANI value calculation are highly variable for genus-level comparisons [52], which raises concerns on the robustness of applying the ANI method to higher taxonomic ranks. To resolve this challenge, analysis of protein sequence divergence among core genes was proposed as a suitable approach [53]. However, while it may be desirable to establish a standardized taxonomy with a defined range of genomic divergence for each taxonomic rank, large variations in the divergence values at a given rank were observed among different taxonomic groups in previous attempts [52–54]. These variations created situations where some families contain higher divergence levels than some orders or lower divergence levels than some genera, even after normalization and a full revision of the current taxonomy [53]. Such situations demonstrated the challenges of establishing a new standardized taxonomy even when the practical issues of transitioning from the current taxonomy are not considered, and perhaps is to be expected given the highly variable evolutionary rates across different lineages [55]. Given these considerations, other aspects of biology (e.g., physiology, ecology, etc) may play more important roles in defining those higher taxonomic ranks.

### Units and Modularity of Molecular Evolution

For evolutionary studies, the levels at which selection and other processes operate on have been a topic that received much attention [56]. For prokaryotes, levels from the entire genome to individual functional domains within genes are of particular interest. Based on our results, all of these levels must be considered to comprehend the complex patterns.

At the whole-genome level, the clear species boundaries based on overall similarity (Figure 1 and Supplementary Figure S1) suggested that the entire genome largely evolves as a single coherent unit. This result is consistent with previous findings that at the global level HGT has very little impact on the reconstruction of organismal phylogeny [57, 58], despite the extensive HGT inferred in bacterial evolution [58–61] and the importance of HGT in adaptation [62–64]. A possible explanation for these seemingly conflicting observations is that most of the acquired genes are lost quickly [63], presumably due to the strong mutational bias towards deletions observed in bacteria [65, 66]. Additionally, acquired genes are subjected to the selection that drives species diversification [8], which is expected to act on all genes in a genome together.

At the level of individual replicons, chromosomes and plasmids certainly have distinct evolutionary histories (Figure 6). Because novel chromosome/plasmid combinations may lead to speciation [50], and the spread of plasmids have important implications on the evolution of virulence [9, 67] and antimicrobial resistance [68], further investigations on the evolution of plasmids and their compatibilities with chromosomes are important [9,69,70]. Additionally, for bacteria with multiple chromosomes, examining the evolution of individual chromosomes may provide novel insights. In the case of *A. tumefaciens*, the multipartite genome was hypothesized to originate from intragenomic gene transfer from the ancestral circular chromosome to a plasmid, followed by linearization of this plasmid to form the secondary chromosome [23]. The secondary chromosome is known to exhibit higher levels of divergence in overall organization, gene content, and sequences [23, 26]. In this regard, it is curious to note that the apparently rapid-evolving T6SS genes are all located on the secondary chromosome, rather than the more conserved primary chromosome. For future studies, it may be interesting to compare the molecular evolution of T6SS and other genes between species with mono- and multi-partite genomes.

At the levels of gene clusters and below, several interesting observations were made based on the loci of the two secretion systems investigated in this work. First, although these two systems may provide some fitness advantages (e.g., T6SS for interbacterial competitions and T4SS for host exploitation), complex patterns of gains and losses were observed (Figures 2 and 6). These patterns suggested that there is not a strong selective pressure to maintain these genes and non-adaptive stochastic processes are important. Furthermore, there is a certain degree of modularity regarding their evolution, such that the presence patterns are all-or-none and no partial cluster was found for the chromosomal T6SS genes or the plasmid-encoded T4SS genes. This is particularly evident for the T6SS genes, as when the main cluster was lost (e.g., G2 and G6), no accessory *vgrG* loci located elsewhere was found (Supplementary Figure S3). Second, for both systems, genes for the structural components are conserved, while those for the effectors and others are not (Figures 2, 5, and 7). This is consistent with the expectation that opposite selective forces may act on these two categories of genes, with purifying selection against changes to preserve the apparatus of a functional secretion system and positive selection for more diverse effectors and other accessory components. Third, modularity that may reflect functional constraints were observed at finer scales. For example, each *vgrG* is linked to its cognate effector/chaperone genes, and each effector is linked to its cognate immunity gene. Such modularity is expected to be maintained by selection, similar to the observations regarding co-transfers of genes involved in associated biochemical pathways [8]. However, the linkage could be broken down by recombination at within- or between-species levels, as evident in the diversity of *vgrG* gene neighborhoods, even for those homologs belonging to the same type (Figure 4). Fourth, at the level of individual genes, the within-genome diversity of *vgrG* homologs (Figure 5) provides further support to the hypothesis that HGT is more important than duplications in driving gene family expansions in bacteria [71]. Finally, at the intra-gene level, within- or between-species recombination may be important in generating novel combinations of domains, thus promoting the diversification of homologs (Figure 3 and Supplementary Figure S6).

Taken together, these observations illustrated the complexity of biological systems. While it is difficult to draw up generalized rules or to estimate the relative importance of each evolutionary process at different levels, it is important to consider and examine these complexities to better understand organisms of interest.

### Conclusions

In summary, by using a group of important bacteria as the study system, this work utilized a strategy of phylogeny-conscious genome sampling for systematic investigations of a species complex. This approach is not only cost-effective but also critical in obtaining an unbiased picture that does not over-emphasize certain subgroups. Additionally, the emphasis on generating and utilizing high-quality assemblies improves the confidence in gene content analysis. With the continuing advancements in sequencing technologies and bioinformatic tools, this emphasis becomes increasingly accessible. For this study system, our examination of biological boundaries at the species level and above improves the understanding of how natural biodiversity is organized. The targeted analysis of those secretion system genes and oncogenic plasmids provides novel insights regarding the key genetic variations involved in the fitness and ecology of these soil-borne phytopathogens that need to compete in complex microbiota and invade plant hosts. Moreover, the multi-level analysis of their genetic diversity from whole-genome to intra-genic domains highlights the complexity of these biological systems. The strategy and findings of this work provide useful guides for future studies of other bacteria.

## Materials and Methods

### Genome Sequencing

A total of 14 strains were acquired from the French International Center for Microbial Resources (CIRM) Collection for Plant-associated Bacteria (CFBP) (Table 1). These include 12 strains that belong to the *A. tumefaciens* species complex and two *A. larrymoorei* strains as the outgroup.

The procedure for whole-genome shotgun sequencing was based on that described in our previous studies [7,72,73]. All bioinformatics tools were used with the default settings unless stated otherwise. Briefly, total genomic DNA was prepared using the Wizard Genomic DNA purification kit (Promega, USA). The Illumina paired-end sequencing libraries were prepared using KAPA LTP Library Preparation Kits (Roche Sequencing, USA) with a targeted insert size of ∼550 bp. The Illumina MiSeq platform was used to generate 300×2 reads with an average coverage of 306-fold per strain (range: 141- to 443-fold). The raw reads were quality trimmed using a Q20 cutoff and used for *de novo* assembly based on Velvet v1.2.10 [74] with the settings “-exp_cov auto-min_contig_lgth 2000-scaffolding no”. The contigs were oriented by mapping to those complete genome assemblies available (Table 1) using MAUVE v2015-02-13 [75]. Due to the difficulties of identifying appropriate reference genomes for several evolutionary branches, four strains (i.e., CFBP5473, CFBP5875, CFBP5877, and CFBP6623) were selected for PacBio long-read sequencing and PacBio HGAP v3 assembly. These PacBio-based assemblies were used as a guide for scaffolding, rather than the finalized results.

To improve the Illumina-based draft assemblies, an iterative process was used to examine the raw reads mapping results and to incorporate gap-filling results based on PCR and Sanger sequencing. This process was repeated until the complete assembly was obtained or the draft assembly could not be improved further. The finalized assemblies were submitted to the National Center for Biotechnology Information (NCBI) and annotated using the Prokaryotic Genome Annotation Pipeline (PGAP) [76].

### Comparative and Evolutionary Analysis

The genomes analyzed are listed in Table 1. The procedures for genome comparisons were based on those described in our previous studies [7,77–79]. Briefly, pairwise genome similarities were calculated using FastANI v1.1 [10]. For comparisons of plasmids, FastANI was executed with the custom settings that reduced fragment length to 1,000 bp and minimum matched fragments to 25. For global alignments of chromosomes and plasmids, the syntenic regions were identified by BLASTN v2.6.0 [34] and visualized using genoPlotR v0.8.9 [80]. For gene content comparison, BLASTP v2.6.0 [34] with e-value cutoff set to 1e^-15^ and OrthoMCL v1.3 [81] were used to infer the homologous gene clusters. The result was converted into a matrix of 35 genomes by 17,058 clusters, with the value in each cell corresponding to the copy number. This matrix was further converted into a Jaccard distance matrix among genomes using the VEGAN package v2.5-6 in R, then processed using the principal coordinates analysis function in the APE package [82] and visualized using ggplot2 v3.3.2 [83]. The hierarchical clustering analysis was performed using PVCLUST v3.4.4 [84].

For phylogenetic analysis, homologous sequences were aligned using MUSCLE v3.8.31 [85] for maximum likelihood inference by PhyML v.3.3.20180214 [86]. The proportion of invariable sites and the gamma distribution parameter were estimated from the data set, the number of substitute rate categories was set to four. The bootstrap supports were estimated based on 1,000 replicates.

### Analysis of the Type VI Secretion System Genes

To identify the T6SS-associated genes, C58 [27,36,37] and other strains [7, 33] that have been characterized experimentally were used as the references. Based on the known T6SS genes in these genomes, homologous genes in other genomes were identified based on the OrthoMCL result. To screen for novel T6SS effector, chaperone, and immunity genes, genes that are located near *vgrG* (i.e., three upstream and ten downstream) were examined manually by using the NCBI conserved domain database (CDD) [87] and the Phyre2 protein fold recognition server [88]; the e-value cutoff was set to 0.01. A few genes with a hit to the DUF4123 domain (i.e., a known T6SS chaperone) but have an e-value above the cutoff were manually added back to the list of T6SS-associated genes (e.g., CFBP5477_RS20350). To confirm the absence of specific T6SS genes, the genome sequences were used as the subjects and the protein sequences of known genes were used as the queries to run TBLASTN searches.

For classification of the *vgrG* homologs, we developed a domain-based scheme. The conserved N-terminal TIGR03361 domain was first identified by the NCBI CDD searches. A global alignment of all homologs was used to determine the exact boundaries of this domain. After this TIGR03361 domain was removed, the remaining C-terminal sequences were processed using MEME v5.1.1 [89] to identify conserved domains that meet these criteria: (1) present in at least two sequences, (2) with zero or one occurrence per sequence, (3) with a size between 30 and 300 a.a., and (4) with an e-value lower than 0.05. The results were manually curated to break down large domains that are composed of smaller domains. Pairwise BLASTP searches were conducted to verify that each domain is unique and no two domains have a BLASTP e-value of lower than 1e-05. For each domain, the consensus sequence was generated using Jalview v2.10.5 [90] and sequence conservation was visualized using WebLogo server v3 [91]. For functional prediction, the consensus sequence of each domain was used to query against NCBI CDD and Phyre2. Additionally, one representative from each subtype of *vgrG* homologs was used for structure modeling using Phyre2 with the “normal” mode. The chain D of PA0091 VgrG1 (PDB identifier: 4MTK) was selected as the template. The predicted structures were visualized using PyMOL v1.2r3pre (Schrödinger, USA).

For the EI gene pairs identified, EI1 through EI11 were named based on the nomenclature proposed previously [7] and novel pairs were named starting from EI12. When only the putative effector (E) or the putative immunity (I) genes were found, those genes were classified in the format of “E??” or “I??”, respectively. For some of the EI pairs that were identified previously based on adjacency to T6SS genes but lacked high-confidence annotation (i.e., EI2, EI3, EI5, and EI8), we chose a more conservative approach and annotated those genes as hypothetical proteins.

### Analysis of the Tumor-inducing Plasmids and Type IV Secretion System Genes

The list of 20 putative pTi sequences analyzed is provided in Table 2. These included all of the 15 complete pTi sequences determined in this study and five representatives from GenBank that are important in *Agrobacterium* research [15]. The pTi typing was performed based on k-mer profile clustering with a reference set of 143 oncogenic plasmids in Rhizobiaceae [9] and a second set that contains > 4,000 Rhizobiaceae plasmids [42]. For T-DNA identification, putative T-DNA borders were identified based on the motif YGRCAGGATATATNNNNNKGTMAWN [92]. Genes involved in opine metabolism [93] and T4SS [94] were identified based on the annotation and homologous gene clustering results produced by OrthoMCL. Additionally, putative T4SS effectors were identified using the T4SEpre tool [95] in EffectiveDB [96] with the minimal score set to 0.8. All protein sequences of pTi-encoded genes were used as the queries.

### Tumorigenesis Assay

Tomato tumorigenesis assays [97] were performed to evaluate the virulence of selected strains. The plants (cultivar Known-You 301) were maintained in growth chambers with a 16-/8-h light/dark regime and a constant temperature of 22°C. Inoculation was performed on three-week old seedlings. Bacterial strains were transferred from stock to 5 mL 523 broth and cultured overnight at 28°C in a shaker incubator (250 rpm), then sub-cultured for four hours prior to inoculation. Bacterial cells were washed and resuspended in 0.9% NaCl solution with a concentration of OD_600_ 0.2. The stem was punctured with a sterilized sewing needle and 5 μL of bacterial suspension was added to the wounding site. The plants were collected three weeks after inoculation and 1-cm stem segments centered at the wounding site were cut for weighing.

## Data Availability

The 14 new genome sequences are available in NCBI under BioProject accessions PRJNA534385-PRJNA534397 and PRJNA534399.

## Author Contribution

Conceptualization: CHK

Funding acquisition: JHC, EML, CHK

Investigation: LC, YCL, MH, MNS, AJW, CFW

Methodology: LC, YCL, MNS, STC, CHK

Project administration: CHK

Supervision: JHC, EML, CHK

Validation: LC, YCL, MH, MNS, AJW

Visualization: LC, YCL

Writing – original draft: LC, YCL, CHK

Writing – review & editing: LC, YCL, AJW, CFW, JHC, EML, CHK

## Acknowledgement

We thank Ai-Ping Chen, Hsin-Ying Chiang, Shu-Jen Chou, Mei-Jane Fang, Ya-Yi Huang, and Wen-Sui Lo for technical assistance. Sophien Kamoun provided helpful comments that improved the writing of this manuscript. The bacterial strains were imported under the permits 103-B-003 and 104-B-002 issued by the Council of Agriculture of Taiwan. The Sanger sequencing service and the Illumina sequencing library preparation service were provided by the Genomic Technology Core (Institute of Plant and Microbial Biology, Academia Sinica). The Illumina MiSeq sequencing service was provided by the Genomics Core (Institute of Molecular Biology, Academia Sinica). The PacBio sequencing and data processing service was provided by Genomics BioSci & Tech. Co. Ltd. (New Taipei City, Taiwan). The Institute of Plant and Microbial Biology (Academia Sinica) and the Department of Botany and Plant Pathology (Oregon State University) provided computing resources.

## Supplementary Materials

**Supplementary Figure S1.**
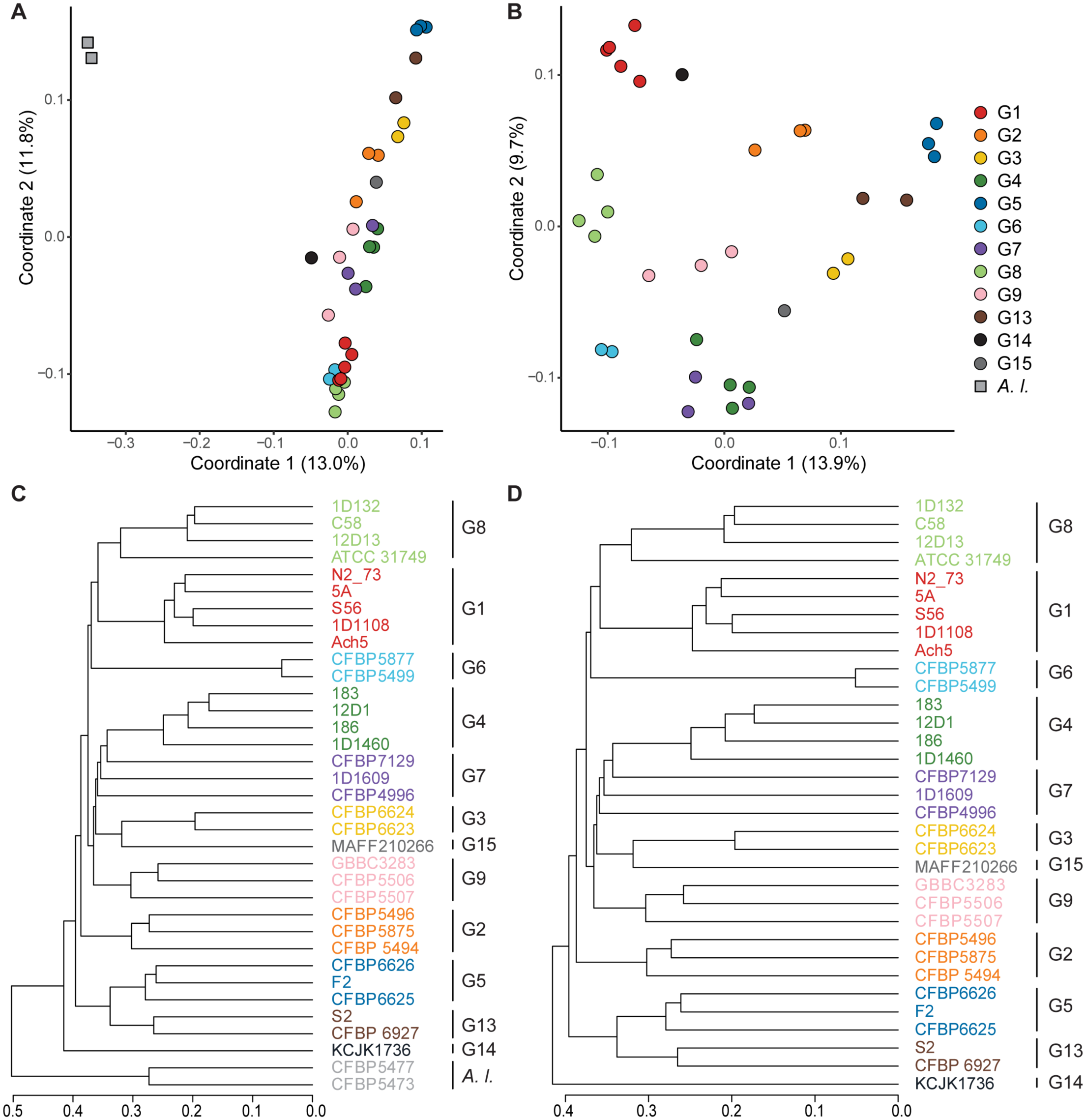
Gene content dissimilarity among the *Agrobacterium* genomes. (A) and (B): principal coordinate analysis with and without the outgroup *A. larrymoorei*, respectively. The % variance explained by each axis is provided in parentheses. (C) and (D): hierarchical clustering with and without the outgroup *A. larrymoorei*, respectively.

**Supplementary Figure S2.**
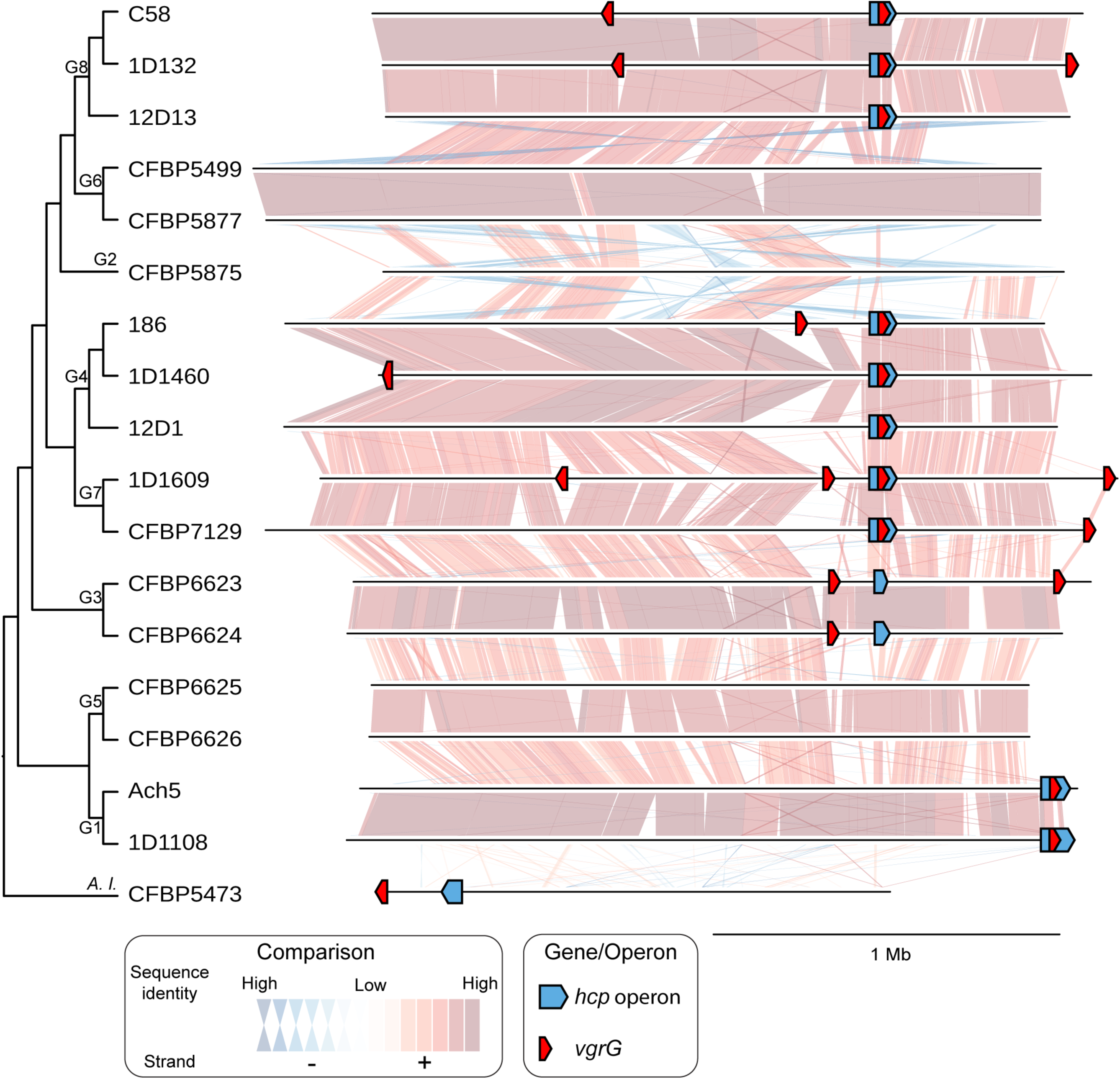
Global alignment of the linear chromosomes. Locations of T6SS-*hcp* operons and *vgrG* homologs are labeled.

**Supplementary Figure S3.**
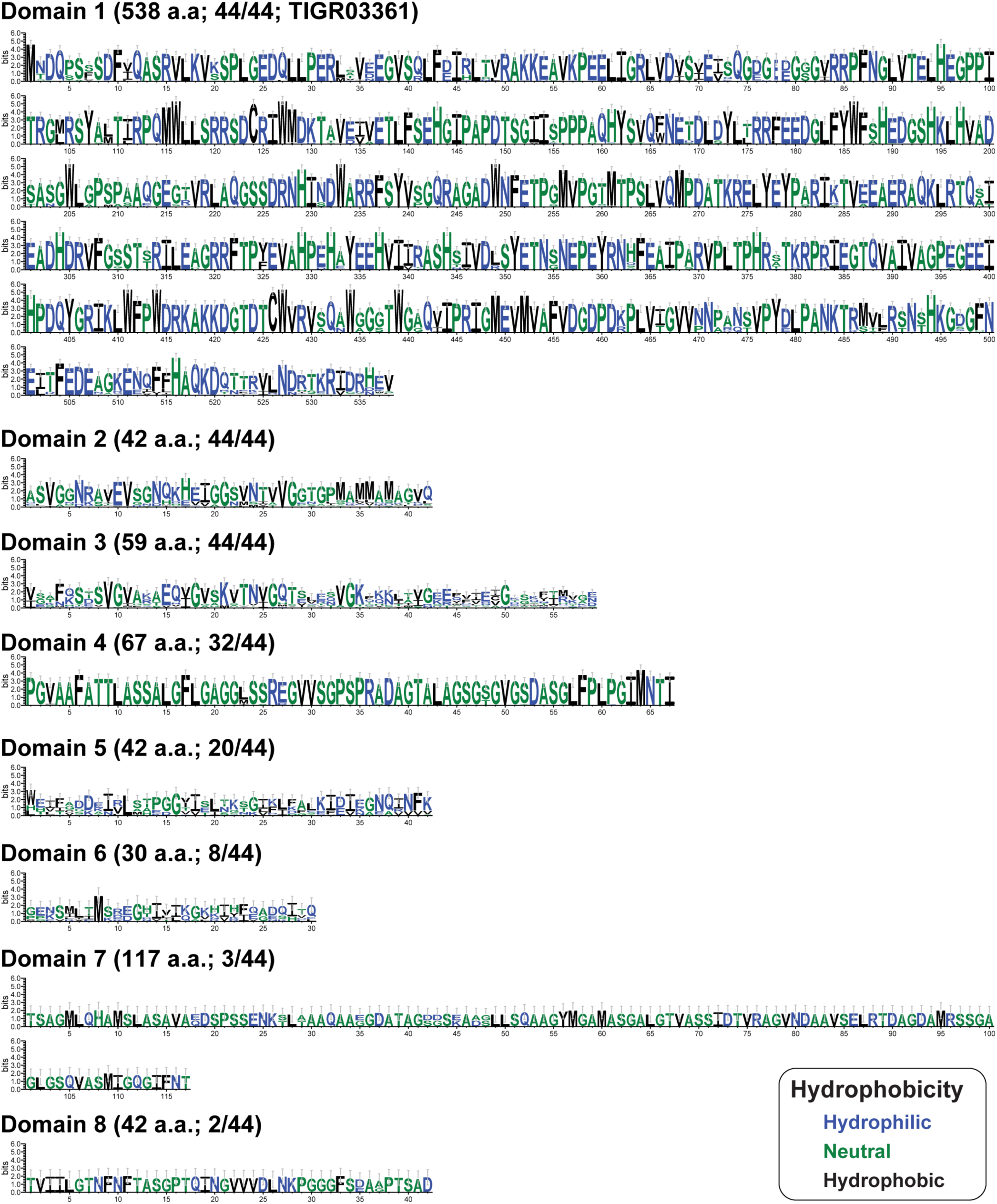
Logo plots of the putative protein domains identified among *vgrG* homologs. For each domain, the length and the number of homologs with the domain is labeled. Domain 1 is the only domain with a corresponding database entry (TIGR03361).

**Supplementary Figure S4.**
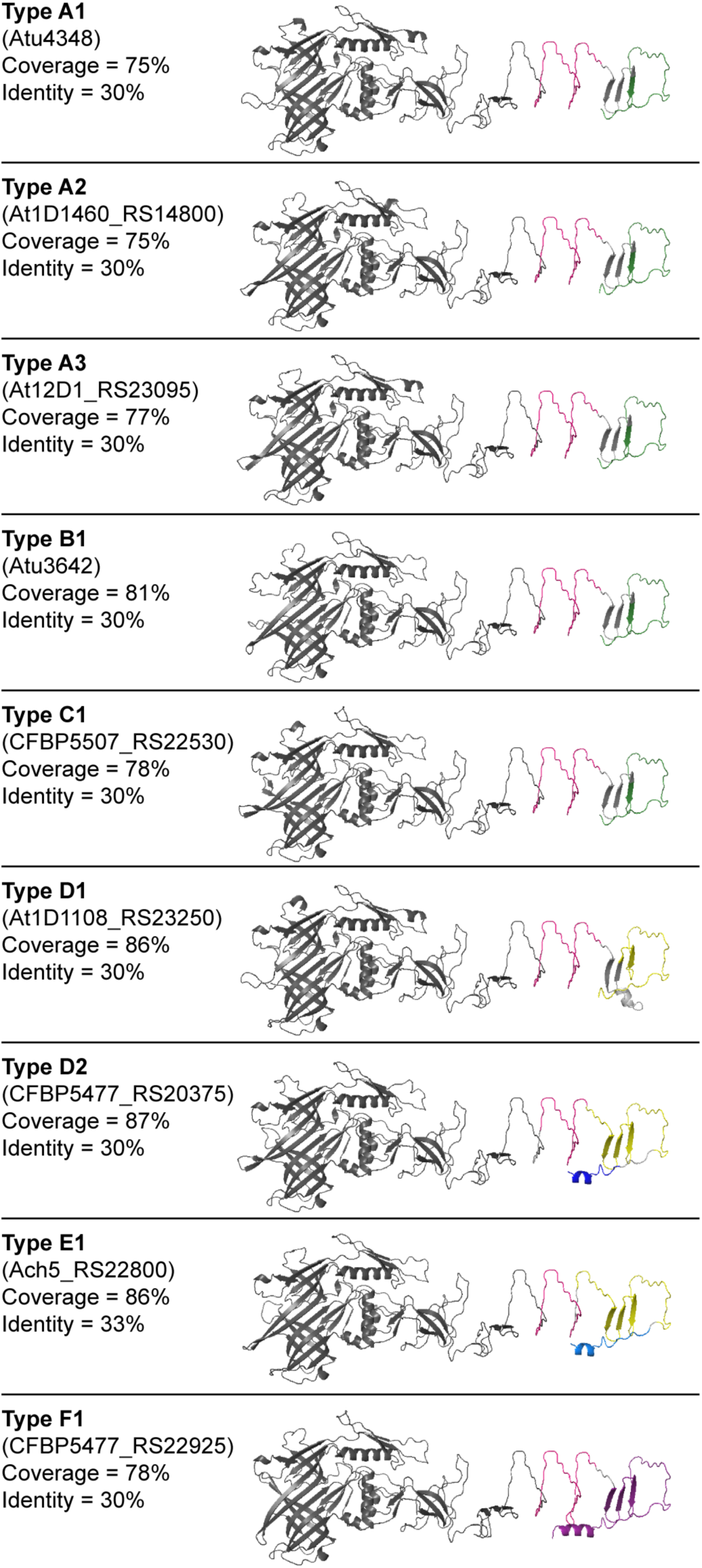
Predicted structures of VgrG homologs. Regions are colored according to the scheme used in the domain analysis (**Figure. 3**). The chain D of PA0091 VgrG1 (PDB identifier: 4MTK) from *Pseudomonas aeruginosa* was selected as the template. The C-terminal parts that could not be confidently inferred are omitted. In all cases, the coverage (i.e., percentage of the sequence included in the structure prediction) are at least 75%, the sequence identity to the template is at least 30% and the confidence score is 100%.

**Supplementary Figure S5.**
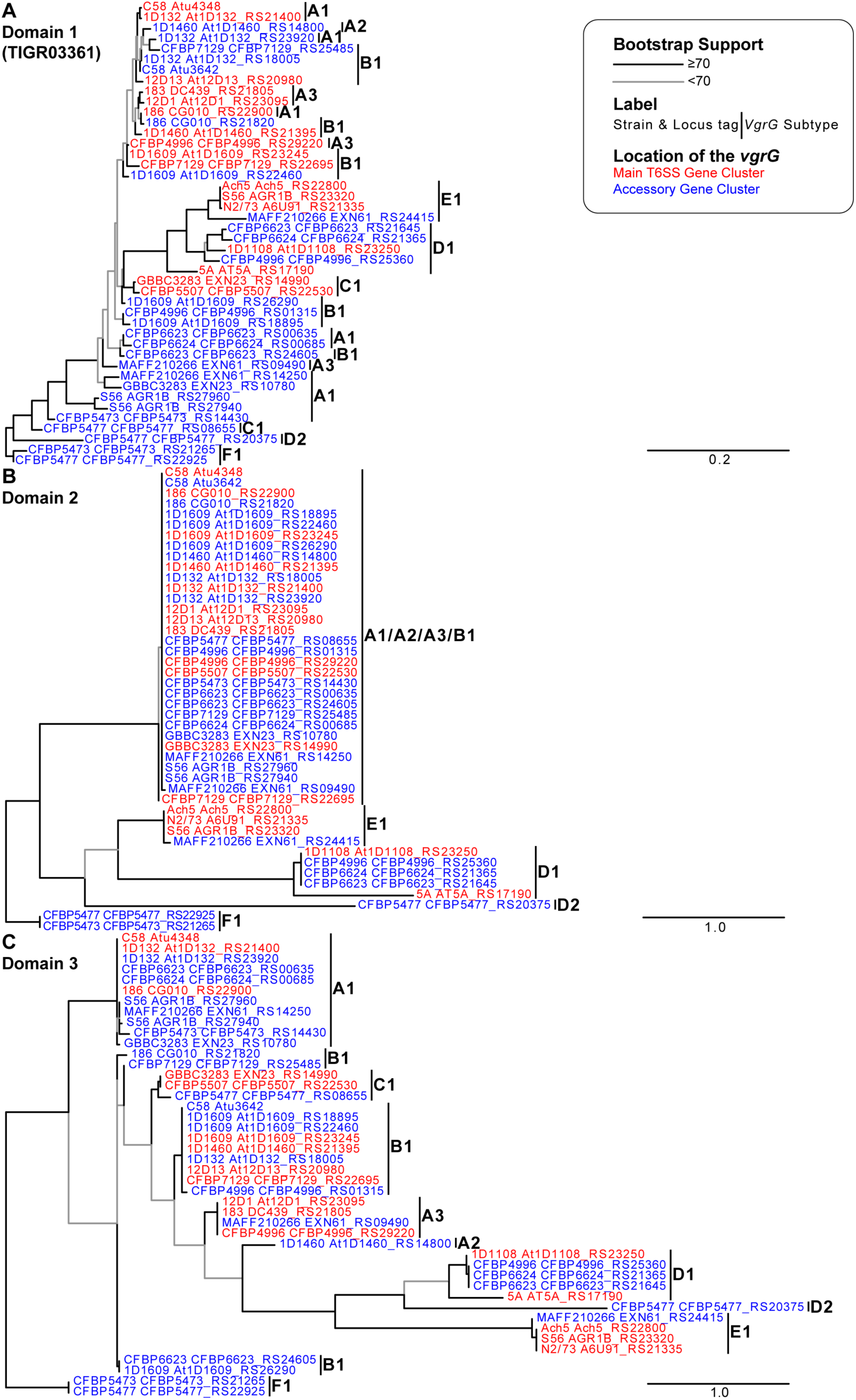
Maximum likelihood phylogenies of *vgrG*-associated domains. (A) Domain 1 (TIGR03361; VI_Rhs_Vgr super family), (B) Domain 2 (unknown function), and (C) Domain 3 (unknown function).

**Supplementary Figure S6.**
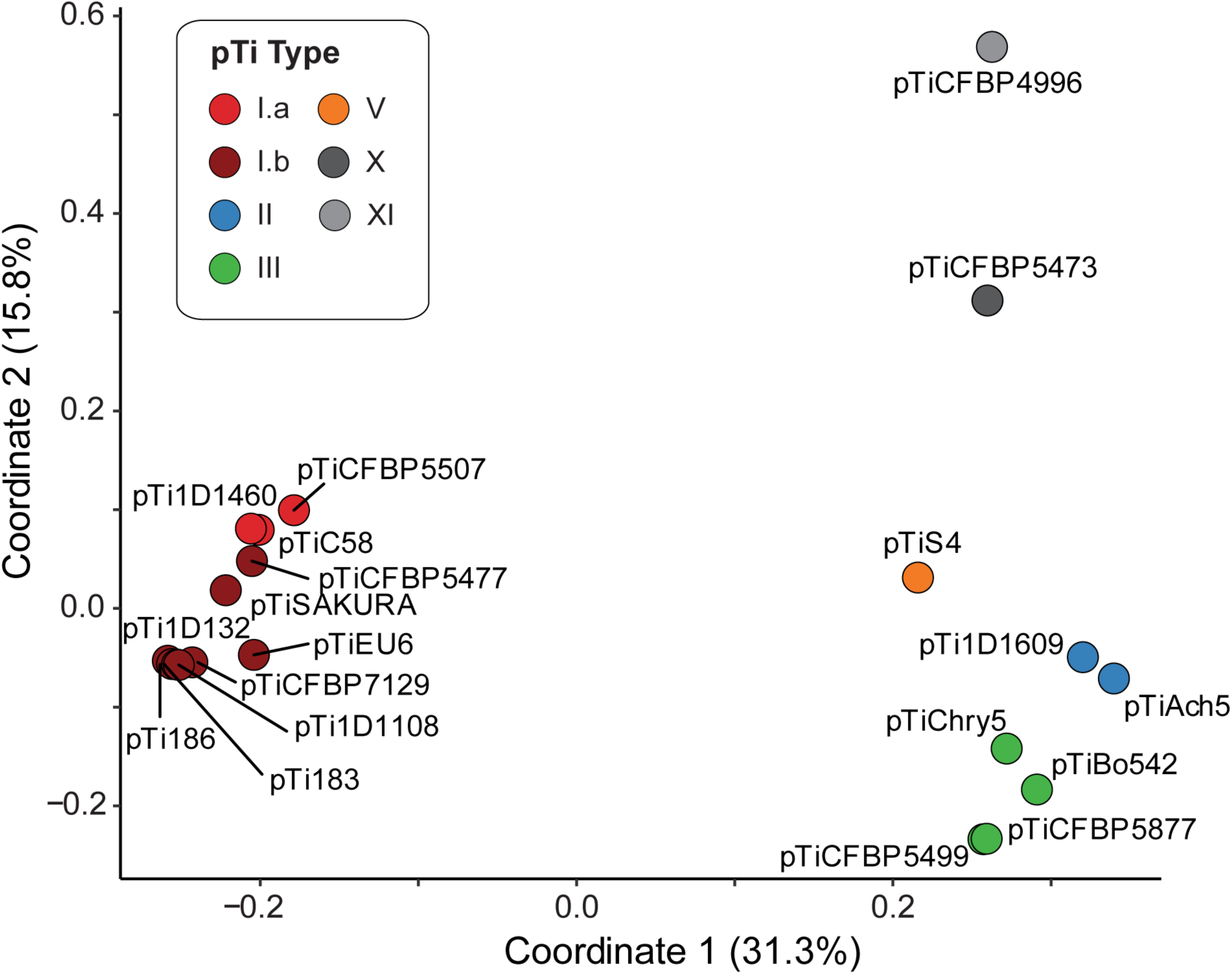
Principal coordinate analysis of gene content among the putative pTi analyzed.

**Supplementary Figure S7.**
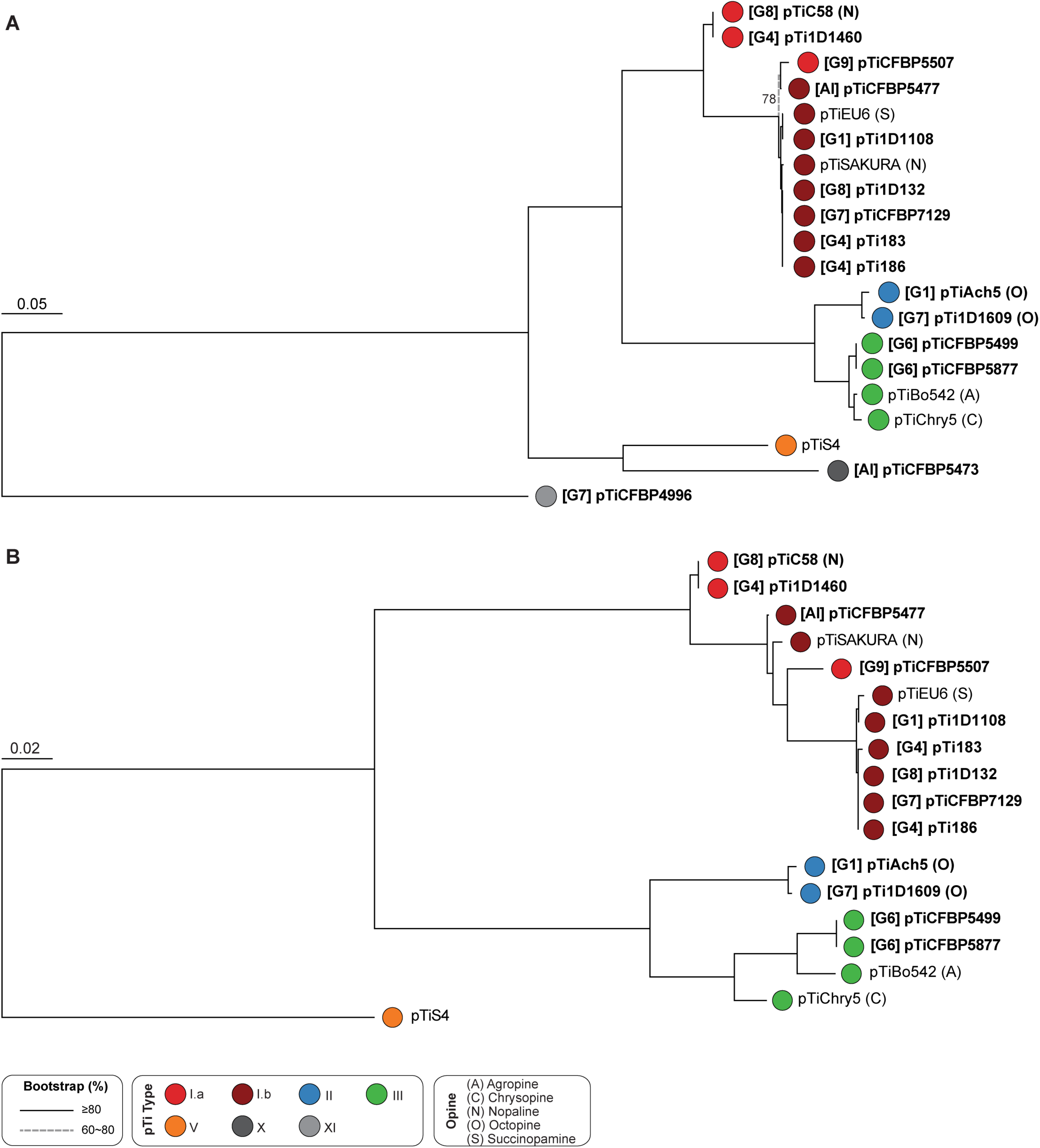
Maximum likelihood phylogeny of pTi based on the concatenated alignment of shared single-copy genes. (A) All of the 20 pTi sequences analyzed; 21 core genes and 8,534 aligned amino acid sites. (B) Excluding the two novel pTi; 40 core genes and 15,473 aligned amino acid sites, all branches received > 80% bootstrap support.

**Supplementary Figure S8.**
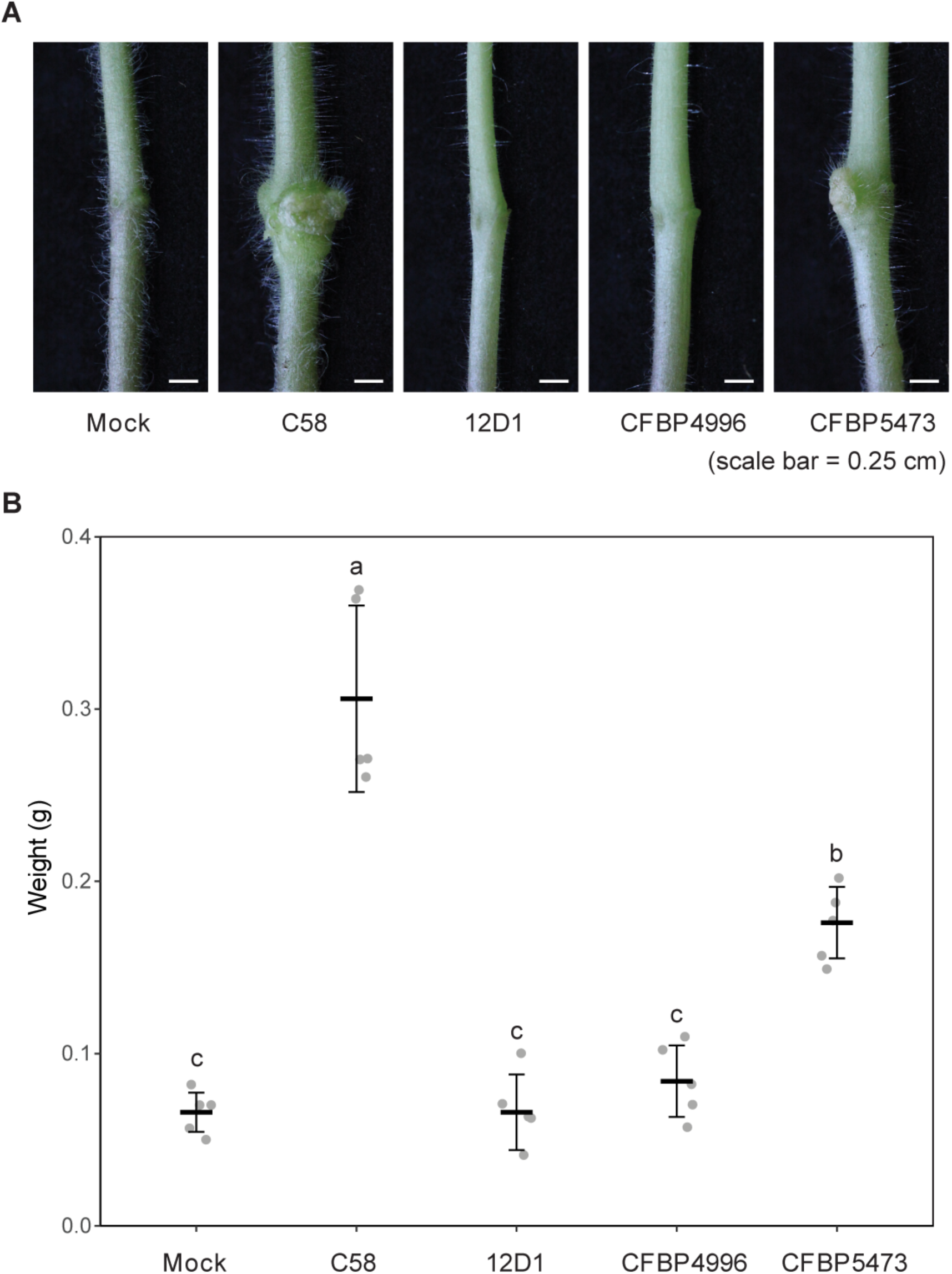
Tomato tumor assay of strains 12D1, CFBP4996, and CFBP5473. Mock was inoculated with sterilized water as a negative control and strain C58 was included as a positive control. Strain 12D1 harbors a plasmid with opine transporter and catabolism genes but lacks *vir* regulon genes and identifiable T-DNA. CFBP4996 and CFBP5473 harbor novel types of putative tumor-inducing plasmids (pTi). (A) Tomato stems at three weeks after inoculation. Scale bar: 0.25 cm. (B) Weight distribution of five biological replicates (1-cm segments of the stem centered at the inoculation site). The letters indicate ANOVA results.

**Supplementary Figure S9.**
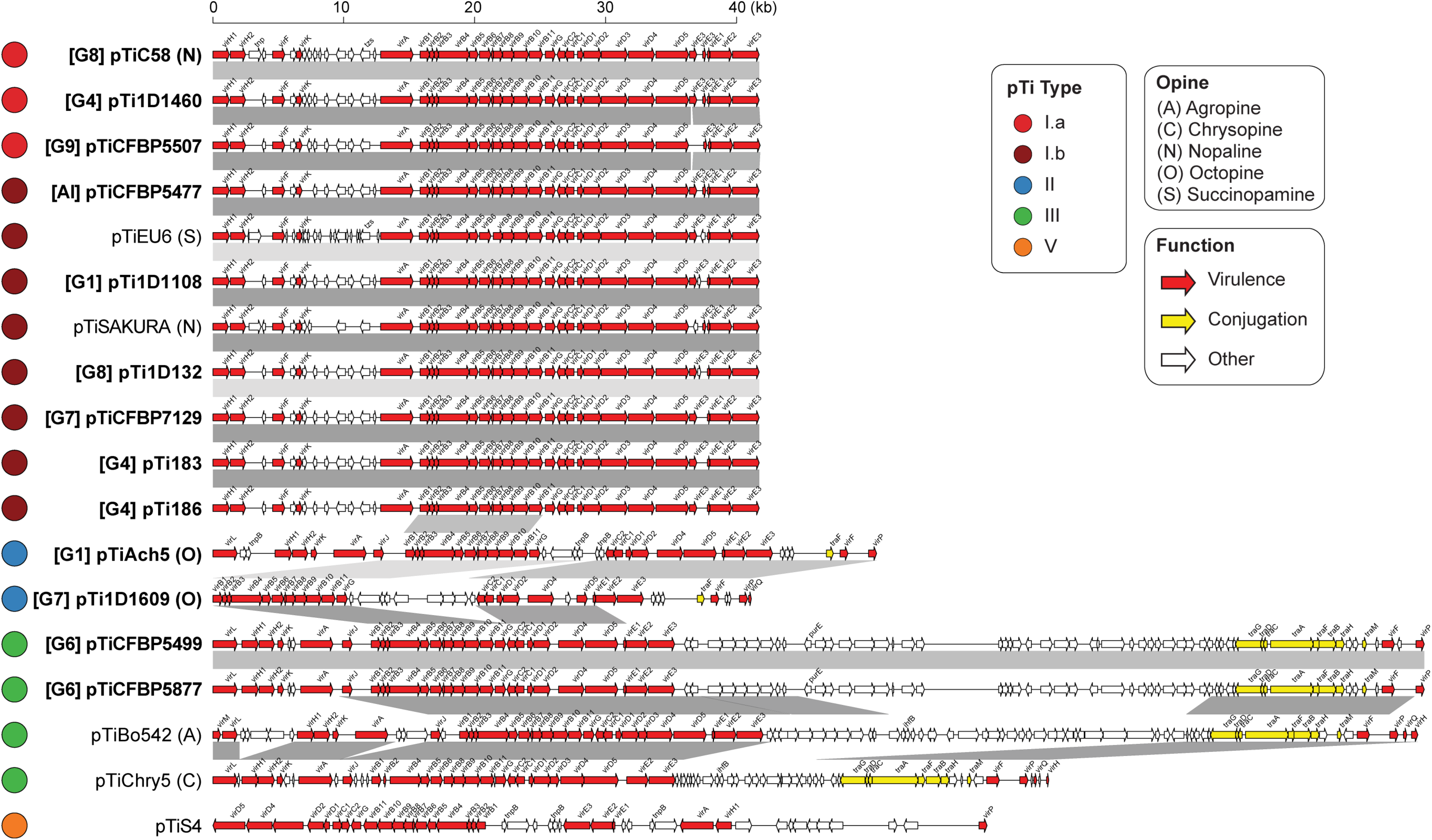
Gene organization of the *vir* regulons on pTi. Syntenic regions are indicated by grey blocks. The virulence (*vir*) genes are highlighted in red, the conjugation (*tra*) genes are highlighted in yellow, and other genes are plotted in white.

**Supplementary Dataset S1.** Detailed information of the analyzed genes. (A) List of *vgrG*-associated genes. Information including genomic location, RefSeq annotation, and domain prediction are included. (B) Locus tags of the *vir* regulon genes on pTi. The *virA*/*J* of 1D1609 are located on another plasmid and are highlighted by “*”. The *virB7* of pTiChry5 and pTiEU6 are unannotated in the GenBank RefSeq records so no locus tag is available but the gene presence was confirmed by BLASTN searches.

